# Spatial dissimilarity analysis in single cell transcriptomics

**DOI:** 10.1101/2025.01.04.631330

**Authors:** Quan Shi, Karsten Kristiansen

## Abstract

The spatial dissimilarity method is a new statistical tool to uncover complex bivariate relationships in single cells and spatial transcriptomics data, thereby addressing unresolved issues such as alternative splicing and allele-specific gene expression. By applying our spatial dissimilarity analysis method on datasets of neurons, tumor, and normal cells, we identified thousands of alternatively spliced genes and instances of allele-specific gene expression. This is particularly evident in the analysis of quiescent and transitioning cell states, where our method reveals the nuanced gene expression dynamics associated with these states. Notably, our findings highlight how allele-specific genetic variants can provide insights into the subclone architecture of normal cells and cancer cells, offering a more comprehensive understanding of cellular heterogeneity and a new insight to understand gene function at specific cell types. The spatial dissimilarity analysis method deepens our comprehension of cellular complexity and gene expression dynamics during cell state transitions.

## Main

Despite recent advancements in single-cell RNA sequencing (scRNA-seq)^1–3^ and spatial transcriptomics^4–6^, most research continues to focus primarily on gene expression, often overlooking other characteristics of sequence data, such as genome coordinate information, alternative splicing (AS)^7,8^, alternative polyadenylation^9^, natural antisense transcripts^10,11^, somatic variants^12^, and allele-specific expression (ASE)^13,14^. While some of these features are well-studied in bulk RNA-seq, their analysis at the single-cell level remains limited due to challenges such as high dropout rates and low gene body coverage in scRNA-seq. Furthermore, existing single-cell methods often rely on predefined cell labels, which can be non-robustness given the different resolution in defining cell types^15,16^.

In parallel, spatial statistics, such as Moran’s I, are widely used in spatial transcriptomics to measure spatial dependency^17^. Furthermore, examining spatial relationships between two genes provides insights into gene regulatory networks and cell-cell interactions^18^. Numerous bivariate spatial correlation methods, including modified *t*-test^19^ and novel statistical measures^20,21^, address spatial dependence issues overlooked by classical Pearson’s correlation^22^. In spatial transcriptomics, these measures are commonly used to assess gene relationships, facilitating clustering and expression module definition in spatial contexts^23,24^. In single-cell analyses, cell-cell similarity graphs replace spatial distances to identify informative genes and modules^24^. Despite these advances, current approaches focus primarily on feature similarity, leaving distinct expression patterns underexplored. For instance, assessing the different expression pattern of an exon with its gene on single cells could uncover critical insights into alternative splicing.

In this article, we introduce the spatial dissimilarity (SD) test, a novel spatial measure to evaluate the spatial autocorrelation of a feature and its dissimilarity relative to a biologically connected feature. The SD test can be applied across 1D cell trajectories, 2D spatial locations, or high-dimensional spaces, ensuring flexibility for diverse datasets. We demonstrate the utility of the SD test with two key applications: AS and ASE. For AS, we examine spatial expression differences between an exon or junction and its associated gene. For ASE, we evaluate spatial expression patterns between one allele and its counterpart allele. Beyond these examples, the SD test can serve as a foundational tool to uncover gene expression dynamics and biological mechanisms across diverse single-cell analyses.

To facilitate implementation, we developed Yano, an R package that integrates seamlessly with the widely used Seurat package^25^. Yano also includes novel features for visualizing UMI depth across cell groups and annotating genetic variants with consequence predictions and information from public databases.

## Results

### Overview of the spatial dissimilarity analysis method

Our goal was to systematically identify spatially dissimilar expression events between molecular features (test features) and their associated features (binding features), requiring only the test feature to exhibit spatial autocorrelation. The analysis consists of five streamlined steps.

Step 1: Annotation and quantification of molecular features using PISA^26^, which processes aligned reads to identify genome location-based features (genes, exons, junctions, exon-excluded reads) and sequence-based entities (SNVs) (Fig. 1b).

**Figure 1.**
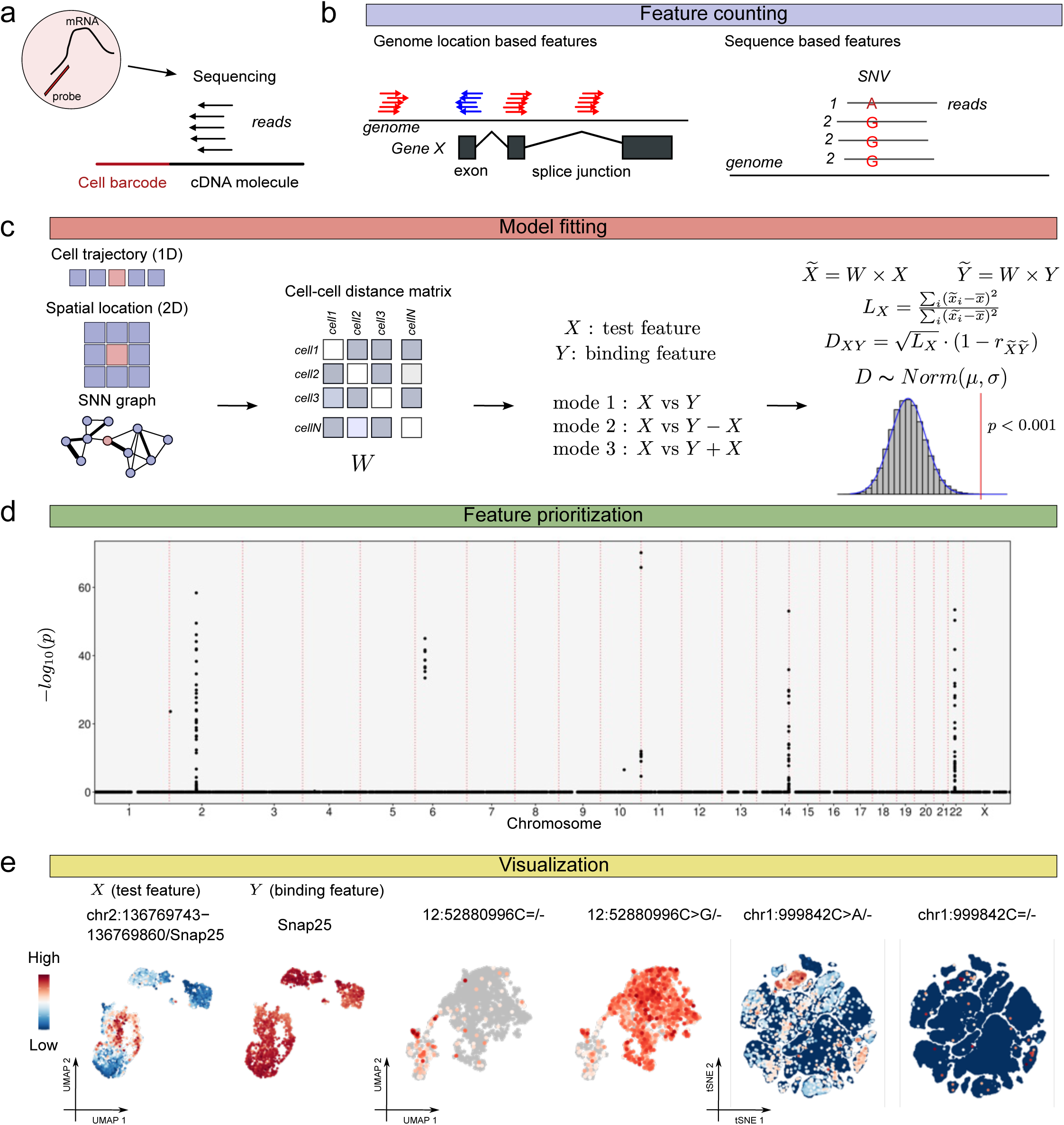
| Overview of spatial dissimilarity analysis method. (a) Simplified pipeline of single-cell RNA sequencing, highlighting key steps such as molecule capture and the sequencing strategy. (b) Counting features derived from alignment reads. This includes genome location-based features like genes, exons, and junctions, as well as sequence-based features such as SNVs. (c) Outlines the process of conducting spatial dissimilarity test. Initially, a cell-cell distance graph is constructed based on either the cells’ positions in a single-cell linear trajectory (1D), their spatial locations (2D), or a SNN graph. Subsequently, cell-cell distance matrix is normalized by column. Then the binding relationship is setup and analysis mode is selected. The D score is calculated with feature counts and weight matrix, and it follows a normal distribution. The P value for D score is calculated to assess the significance of spatial dissimilarity. (d) Genomic locations of features and P values plot on a Manhattan plot. The P value is transformed using a negative log10 scale for visualization. The genomic locations sorted by chromosome and coordinates. (e) Visualizing features on reduction map. (Left) Examples of an exon and its gene on UMAP. (Middle) Allele expression at chromosome 12 position 52880996 on UMAP. (Right) Allele expression at chromosome 1 position 999842 on tSNE.

Step 2: Construction of a cell-to-cell distance/similarity matrix using cell coordinates along developmental trajectories^27^, spatial locations^17^, or shared nearest neighbors (SNN) graphs^25,28^ build on high-dimensional PCA or integrated spaces (e.g., Harmony^29^). The cell-cell distance/similarity matrix is then transformed into a cell-cell weight matrix, which is column-normalized and is directly used to compute Moran’s I^30^. Only features with Moran’s I > 0 and an adjusted P value < 0.01 (Benjamini-Hochberg^31^) proceed to downstream analysis.

Step 3: Establishing the association between test and binding features based on the biological question. For example, an exon can be paired with its corresponding gene to study alternative splicing or one allele with its counterpart to investigate ASE. Three operational modes define the input for binding features: (1) Mode 1: Direct use of binding feature counts after normalization (usually load from ‘data’ layer of Seurat object^25^). (2) Mode 2: Binding counts subtracted by test feature counts before normalization. (3) Mode 3: Binding and test feature counts summed before normalization.

Step 4: Performing the SD test to identify significant feature pairs. We introduce the D score, which integrates the spatial autocorrelation of the test feature (L) and the dissimilarity between the test and binding features (1 - *r*, where *r* is the Pearson correlation of spatial lags) (Fig. 1c). A permutation-based method establishes the distribution of D scores, revealing a normal distribution (Supplementary Fig. 1). Yano software samples the test feature’s distribution 100 times while keeping the binding feature fixed, calculating P values using a one-tailed *t* statistic. High D scores indicate both strong spatial autocorrelation (large L) and low correlation (*r*), reflecting significant spatial dissimilarity.

Step 5: Prioritization of features based on P values and genomic locations (Fig. 1d). Clustered P values in nearby genomic regions offer stronger evidence for significant events. Results are visualized using dimensionality reduction maps and track plots. For example, distinct spatial dissimilarity of an exon at chromosome 2 (positions 136769743-136769860) and its gene *Snap25* indicates an alternative splicing event (Fig. 1e, left). Similarly, contrasting expression patterns of alleles C and G at chromosome 12 suggest loss of allele-specific expression in specific cells (Fig. 1e, middle). Importantly, the SD test evaluates the spatial dependency of the test feature without assumptions about the binding feature’s distribution, enabling the identification of loci with informative expressed alleles and low-expressed counterparts caused by somatic mutations or allele-selective gene repression (Fig. 1e, right).

### Spatial dissimilarity analysis reveals alternative splicing features in mouse primary motor cortex

We applied our SD analysis to four published datasets generated using Smart-seq2^1^, 10X Genomics 3’ v3, and 5’ kits, covering the most commonly used library structures. We analyzed 33,257 neurons from six 10X Genomics 3’ v3 libraries and 4,135 neurons from a Smart-seq2 library, both derived from the mouse primary motor cortex^32^. Cells were clustered and integrated at the gene level using Seurat^25^ and Harmony^29^, with annotations adapted from the original dataset^32^. Exon and junction counts were annotated using PISA, resulting in 62,422 junctions (∼2x genes) and 584,173 exons for 10X libraries, and 166,257 junctions (∼7x genes) and 304,217 exons for Smart-seq2.

We calculated D scores and P values for exon-gene and junction-gene pairs and visualized the results on Manhattan plots (Fig. 2b, c), detecting 1,317 features in 10X data and 2,100 in Smart-seq2 (adjusted P < 1e-10). Splitting the 10X cells into two groups (16,628 cells each) and re-running the test yielded highly consistent results (*r* = 0.97) (Fig. 2d), confirming the robustness of our method.

**Figure 2.**
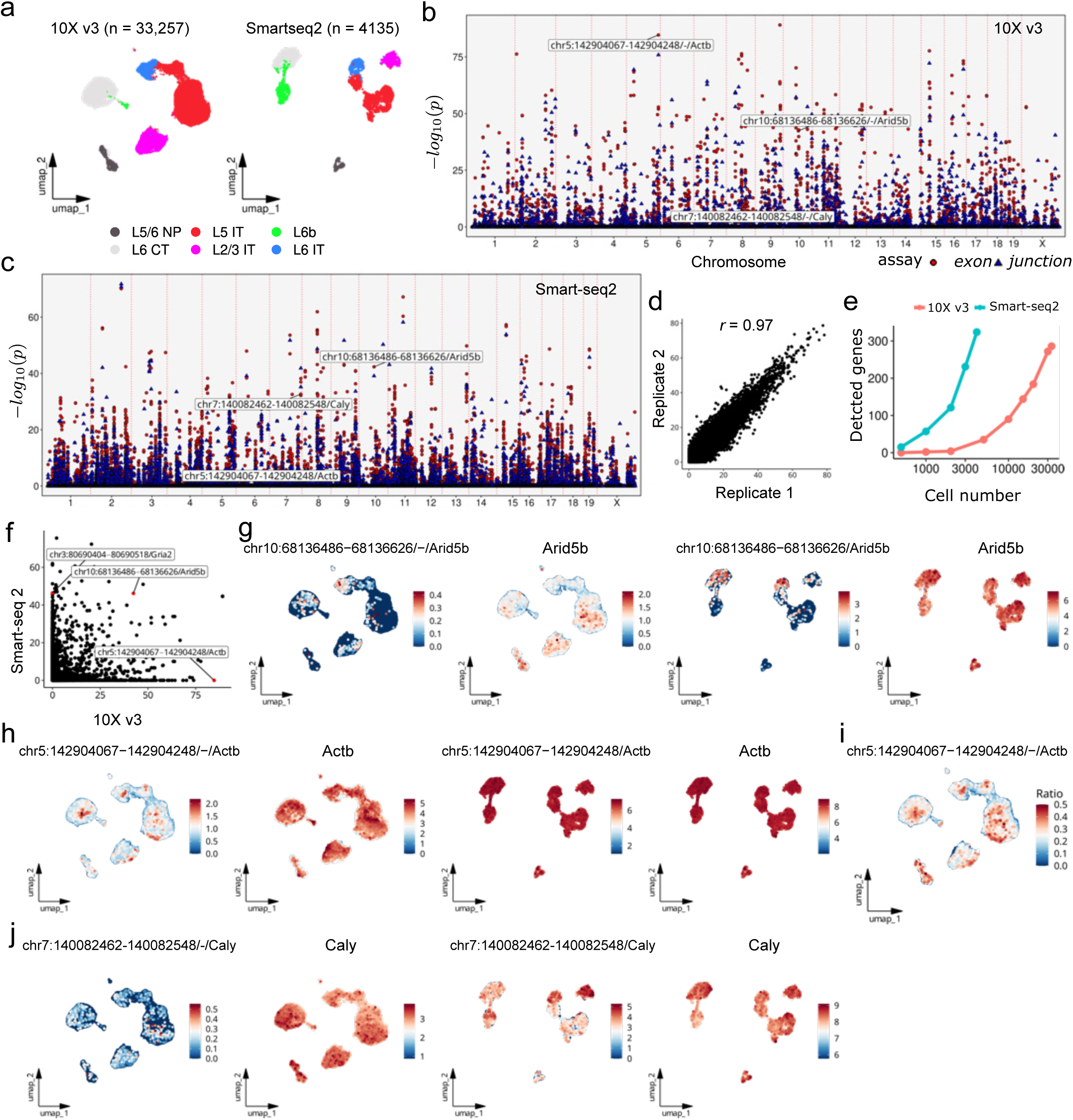
| Spatial dissimilarity analysis reveals alternative splicing features in mouse MOp. (a) UMAP visualization of MOp from the 10X Genomics v3 library (left) and the Smart-seq2 library (right), with cells color-coded by cell types. CT, corticothalamic; IT, intratelencephalically projecting; NP, near-projecting. (b) Manhattan plot illustrating the results of the spatial dissimilarity test between exon/junction and gene pairs for 10X Genomics v3 data. Points were color-coded and shape-coded by the assay types and sorted by genomic coordinates along the x-axis. P value of each event was transformed using a negative log10 scale for visualization. (c) Like (b), this Manhattan plot focuses on the spatial dissimilarity test between exon/junction and gene pairs for Smart-seq2 data. Points are differentiated by color and shape based on the assay type and sorted by genomic coordinates along the x-axis. P value of each event was transformed using a negative log10 scale for visualization. (d) Correlation plot of adjusted P values from two groups of cells. These P values were derived from 10X Genomics v3 data, where cells were randomly divided into two groups, and the SD test was performed separately for each group. The P values were transformed using a negative log10 scale for visualization. Each point represents a feature, illustrating the consistency of adjusted P values between the two groups. (e) Downsampling analysis of 10X Genomics v3 and Smart-seq2 data. Detected genes represent those with at least one event having an adjusted P value < 1e-10. (f) Correlation plot of adjusted P values for exon and gene pairs from 10X Genomics v3 and Smart-seq2 data. P values along the x-axis and y-axis were transformed using a negative log10 scale for visualization. Strand information in exon names from 10X data was removed to enable comparison with Smart-seq2 data. (g) UMAP plots showing the expression of exon chr10:68136486-68136626 and the gene *Arid5b* in 10X Genomics v3 data (left) and Smart-seq2 data (right). Expression levels are normalized on a log scale and adjusted for library size. Note that due to differences in library sizes between assays, expression values are not directly comparable across plots. (h) UMAP plots showing the normalized expression of exon chr5:142904067-142904248 and the gene *Actb* in 10X Genomics v3 data (left) and Smart-seq2 data (right). (i) UMAP plots showing the ratio of raw counts for exon chr5:142904067-142904248 to the gene *Actb* across individual cells. (j) UMAP plots showing the normalized expression of exon chr7:1400882462-149982548 and the gene *Caly* in 10X Genomics v3 data (left) and Smart-seq2 data (right).

To compare full-length sequencing (Smart-seq2) and 3’ biased sequencing (10X) data, we downsampled cells and performed the SD test. With equal cell numbers, 10X data detected fewer genes due to lower gene body coverage (Fig. 2e). The number of detected genes increased with more cells, though saturation was not observed, even with >30,000 cells in the 10X data.

We compared prioritized exons between the two libraries and found 142 exons shared (adjusted P < 1e-10), while 684 were uniquely detected by Smart-seq2 and 574 by 10X (adjusted P < 1e-10 for Smart-seq2, adjusted P < 1e-2 for 10X) (Fig. 2f). Notably, exons unique to 10X often exhibited spatially restricted expression patterns. For example, exon chr5:142904067-142904248 showed spatial expression in nearby cells, whereas its gene *Actb* was globally expressed in 10X data (Fig. 2h, left). In Smart-seq2, this exon’s expression aligned with the gene’s pattern (Fig. 2h, right). Plotting the exon-to-gene count ratio confirmed the distinct spatial pattern in 10X data, ruling out low gene body coverage as a cause (Fig. 2i). Conversely, exon chr7:140082462-140082548, detected only by Smart-seq2, likely reflected low gene body coverage (Fig. 2j and Supplementary Fig. 2).

To systematically summarize alternative expression events, we co-clustered exon and gene counts in Smart-seq2 data and generated a heatmap, revealing complex alternative splicing patterns (Supplementary Fig. 3). Although Smart-seq2 excels at detecting alternative splicing due to its full-length coverage, its non-strand-specific nature can complicate analysis. For instance, *Stk16* overlaps *Tuba4a* on the opposite strand. The high expression of *Tuba4a* causes exons of *Stk16* to show low P values, introducing false positives. In contrast, the strand-specific 10X data clearly distinguishes these genes based on read orientation (Supplementary Fig. 4).

### Benchmarking of the spatial dissimilarity test method against cluster-based methods

While established methods for single-cell AS analysis typically focus on detecting differential AS events between cell groups, they may overlook ‘local’ events within specific subpopulations. To benchmark our SD test, we selected two clusters (L5 IT and L6 IT) from the Smart-seq2 dataset. Rerunning the SD test on these clusters revealed novel AS events missed in the global analysis (Supplementary 5a, b). For example, exon chr9:78478449-78478858 of *Eef1a1* displayed a distinct local expression pattern that was diluted when analyzing all cells (Supplementary Fig. 5c).

We compared our SD test with DEXSeq^33^ and BRIE2^34^, tools use either feature counts or PSI values as input and represent the most commonly used approaches for single-cell AS analysis. DEXSeq detected 2,372 exons (adjusted P < 1e-10), whereas our SD test identified 400 exons. DEXSeq’s sensitivity highlights exon usage differences but struggles to distinguish low gene expression (as in *Ly6e*; Fig. 3d and Supplementary Fig. 5d) from true AS events. We noted exon chr12:4799657-40799750 displayed expression pattern matching *Dock4*, leading the SD test to deprioritize it (Fig. 3e, Supplementary Fig. 6). Conversely, 224 exons were exclusively detected by the SD test (adjusted P < 1e-10), characterized by pronounced differences between exon and gene expression patterns (Fig. 3f, g).

**Figure 3.**
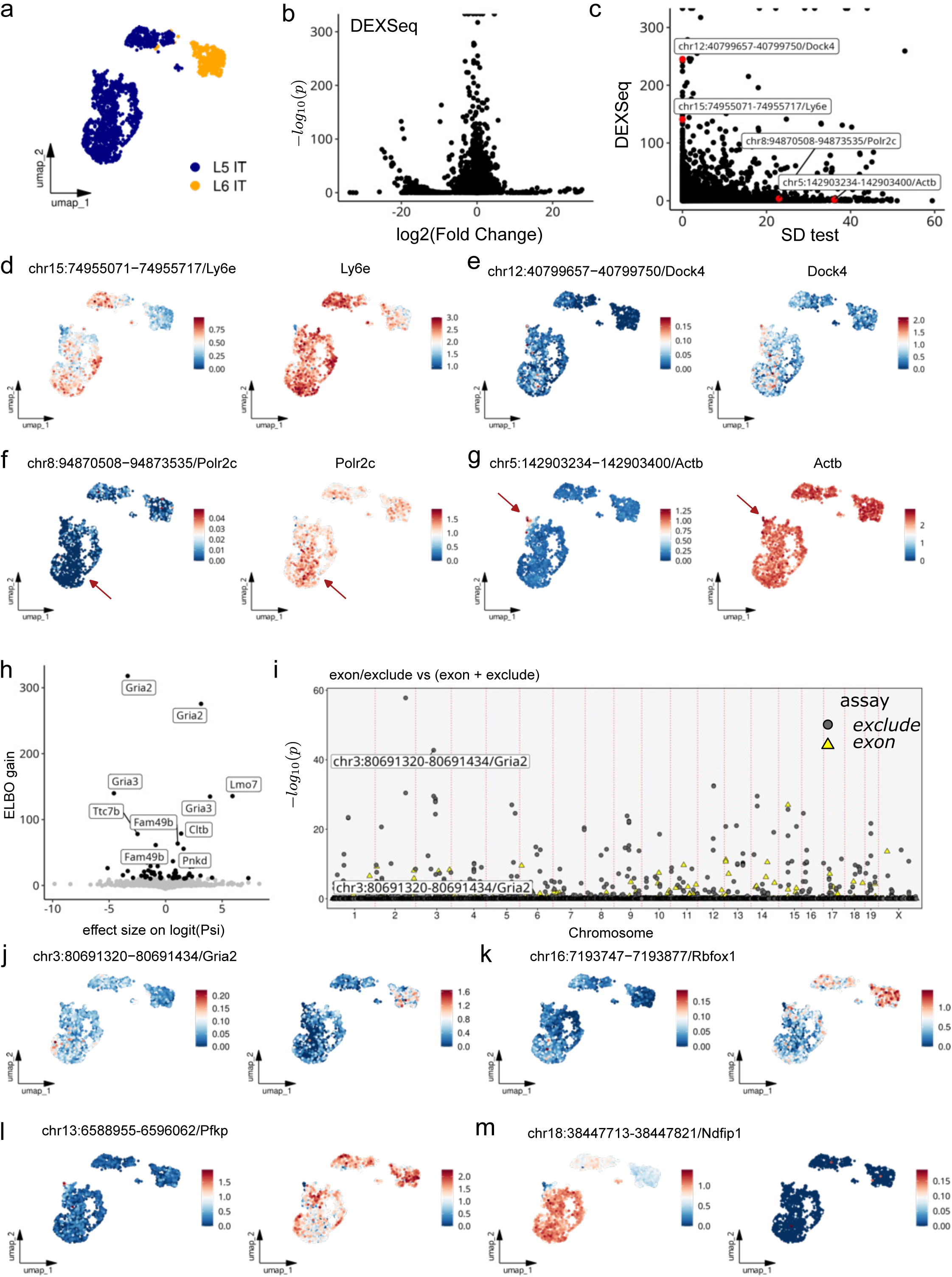
| Benchmarking of the spatial dissimilarity test method against cluster-based methods. (a) UMAP plot of L5 IT and L6 IT cells from Smart-seq2 data, with cells colored by cell type. (b) Volcano plot of DEXSeq results. The x-axis represents the log2 fold change between L5 IT and L6 IT, while the y-axis shows P values transformed using a negative log10 scale for visualization. (c) Comparison of same exon analyzed with DEXSeq and the spatial dissimilarity test results. P values are negative log10 scaled. (d) UMAP plots showing the expression of exon chr15:74955071-74955717 (left) and the gene *Ly6e* (right). Expression levels are normalized on a log scale and adjusted for library size. Note that due to differences in library sizes between gene assay and exon assay, expression values are not directly comparable across plots. (e) UMAP plots showing the normalized expression of exon chr12:40799657-40799750 (left) and the gene *Dock4* (right). (f) UMAP plots showing the normalized expression of exon chr8:94870508-94873535 (left) and the gene *Polr2c* (right). The arrow highlights cells exhibiting an inverse expression pattern between the exon and the gene. (g) UMAP plots showing the normalized expression of exon chr5:142903234-142903400 (left) and the gene *Actb* (right). The arrow highlights cells exhibiting an inverse expression pattern between the exon and the gene. (h) Volcano plot of BRIE2 results. The x-axis represents the effect size difference of PSI between L5 IT and L6 IT, while the y-axis shows the ELBO gain. Exons with low ELBO scores (<10) are colored in grey, while other exons are colored in black. (i) Manhattan plot illustrating the results of the spatial dissimilarity test between exon assay and exon-excluded assay (and vice versa) in Mode 3. Points are color- and shape-coded based on assay type and sorted by genomic coordinates along the x-axis. P values for each event were transformed using a negative log10 scale for visualization. (j) UMAP plots showing the normalized expression of exon chr3:80691320-80691434 (left) and exon excluded reads for the same exon (right). (k) UMAP plots showing the normalized expression of exon chr16:7193747-7193877 (left) and exon excluded reads for the same exon (right). (l) UMAP plots showing the normalized expression of exon chr13:6588955 - 6596062 (left) and exon excluded reads for the same exon (right). (m) UMAP plots showing the normalized expression of exon chr18:38447723-38447821 (left) and exon excluded reads for the same exon (right).

For BRIE2, which prioritizes AS events using the evidence lower bound (ELBO) score^34^, we compared results using Mode 3 (summing exon-included and excluded reads as binding features) and Mode 1. Mode 3 highlighted 46 genes with significant differential patterns (adjusted P < 1e-10), whereas the more sensitive Mode 1 identified 1,985 genes (Supplementary Fig. 7a). As expect, all events in Mode 3 were also detected in Mode 1. Shared results between BRIE2 and Mode 3 included *Gria2* and *Rbfox1* (Fig. 3j, k). Five genes detected by BRIE2 (ELBO > 30) were not identified by our SD test with Mode 3 but can be detected with Mode 1. Unlike BRIE2, which outputs gene-level scores without pinpointing specific splicing sites, the SD test provides site-specific resolution. Upon inspection, we found that minor differences were observed in spliced reads for these genes, though these differences were not substantial (Supplementary Fig. 7b-d). Additionally, we noted that the genes exclusively detected with our SD test often were influenced by alternative expression events within L5 IT (Fig. 3l). Furthermore, high L scores for the test feature combined with sparse expression of the binding feature can also result in high D scores in Mode 1 (Fig. 3m).

Methods like DEXSeq and BRIE2, which rely on linear regression and cluster-based assumptions, exhibit high sensitivity but often introduce false positives or fail to account for within-group expression patterns. In contrast, the SD test balances sensitivity and specificity by directly comparing expression patterns between test and binding features. Importantly, the SD test does not require predefined cell groups, enabling the discovery of hidden cell populations and supporting flexible feature relationships to investigate diverse alternative expression events. Despite these advantages, SD test may lose sensitivity when test and binding features exhibit identical patterns, as high correlation (r) reduces the D score. To address this, we integrated DEXSeq into our pipeline via the RunDEXSeq function for group-based AS testing. Results from RunDEXSeq should always be cross-validated to distinguish with false positive events.

### Spatial dissimilarity analysis of SNVs reveals allele specific gene expression in squamous cell carcinoma

We analyzed 3,464 cells from tumor and paired normal tissues of a human cutaneous squamous cell carcinoma (cSCC) sample^35^(Fig. 4a). Genetic variants were collated from whole exome sequencing (WES) and scRNA-seq data, merged into a VCF file, and post-annotated to quantify expression levels for all alleles at each locus (Methods). To accommodate both single nucleotide variants (SNVs) and reference alleles, we defined expressed allele-specific tags (EATs) to represent alleles throughout this study. Unlike gene expression, directly derived from alignment reads, locus expression was defined as the summed counts of all EATs at a locus. Using the SD test in Mode 2 (comparing other alleles at the same locus), we identified 229,507 EATs across 194,018 loci, with strand-specific loci analyzed independently. Loci with fewer than two EATs were excluded, leaving 31,876 loci for analysis, of which 2,240 showed significant allele-specific expression (adjusted P < 1e-10) (Fig. 4c).

**Figure 4.**
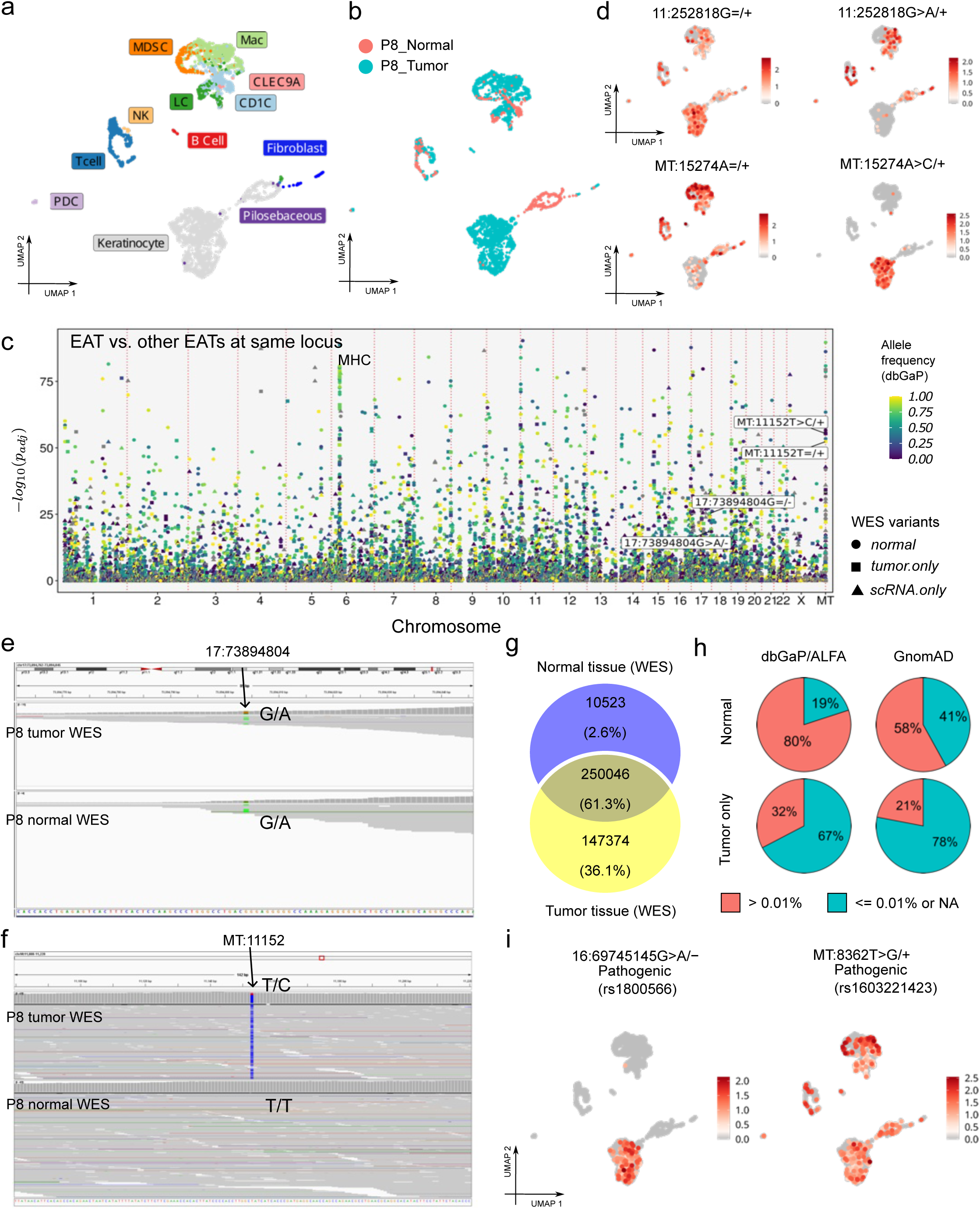
| Spatial dissimilarity analysis of EATs reveals allele-specific gene expression in cSCC. (a) UMAP visualization of cSCC cells, each color-coded according to identified cell clusters. Myeloid-derived suppressor cells, MDSC; Natural killer, NK; Macrophage, Mac; CD1C+ dendritic cells, CD1C; CLEC9A+ dendritic cells, CLEC9A; Langerhans cells, LC. (b) UMAP visualization of cSCC cells colored by tissue type. (c) Manhattan plot illustrating the results of the spatial dissimilarity test between EATs and other EATs at the same locus. Points are color-coded by AF from the dbGaP/ALFA database, shaped by variant origin, and sorted by genomic coordinates along the x-axis. P values for each event were transformed using a negative log10 scale for visualization. (d) UMAP plots showing the expression of EAT 11:252818 allele G and allele A (top) and EAT MT:15274 allele A and allele C (bottom). Expression levels are normalized on a log scale and adjusted for library size. As both EATs at each locus are derived from the same assay, their expression values are directly comparable. (e) IGV plot showing reads from WES data of tumor and normal tissues overlapping with chromosome 17, position 73894804. (f) IGV plot showing reads from WES data of tumor and normal tissues overlapping with mitochondria, position 11152. (g) Venn diagram showing the genetic variants detected in WES data from normal tissues and tumor tissues. (h) Pie charts representing the proportion of low AF somatic mutations (detected in tumor tissues only) across different human population AF databases: dbGaP/ALFA databse (left) and the Genome Aggregation database (GnomAD) (right). (i) UMAP plots showing the expression of two known pathogenic mutations: rs18800566 allele A (left) and rs1603221423 allele G (right), across cells.

Cell-type-specific EATs were detected; for instance, at locus 11:252818, allele G was expressed in myeloid cells and keratinocytes across all samples, while allele A was lost in epithelial tumor cells (Fig. 4d). Conversely, allele C at MT:15274 emerged in keratinocytes specifically within the tumor (Fig. 4d). Further analysis of WES data confirmed that both alleles (G and A) at chromosome 17:73894804 were present in normal and tumor DNA, indicating allele A silencing in keratinocytes within the tumor (Fig. 4e). In contrast, allele C at mitochondrial position 11152 resulted from a somatic mutation (Fig. 4f).

Around 36% of tumor mutations were absent in paired normal cells (Fig. 4g). Annotation with GnomAD^36^ and dbGAP/ALFA (www.ncbi.nlm.nih.gov/snp/docs/gsr/alfa/) showed that these mutations often had lower allele frequency (AF) (Fig. 4), suggesting that rare variants (AF < 0.0001) or unreported alleles are more likely to be somatic mutations. Further annotation with the ClinVar^37^ database identified pathogenic mutations expressed in epithelial and myeloid cells, potentially acquired during cell dividing (Fig. 4i).

In summary, our SD test, combined with genetic variant annotation, successfully identified ASE events and informative somatic mutations in cancer cells. This approach offers significant potential to enhance our understanding of cancer etiology and tumor evolution.

### Identification of squamous cell carcinoma subclones with alternatively expressed EATs

We explored the utility of alternative EAT expression patterns in identifying tumor subclones within a tumor sample. Traditional gene expression analyses rely on highly variable genes (HVGs) for PCA and cell clustering, which can obscure ASE signals. Instead, we re-clustered keratinocytes using PCA and UMAP analyses based on highly variable EATs (Fig. 5a).

**Figure 5.**
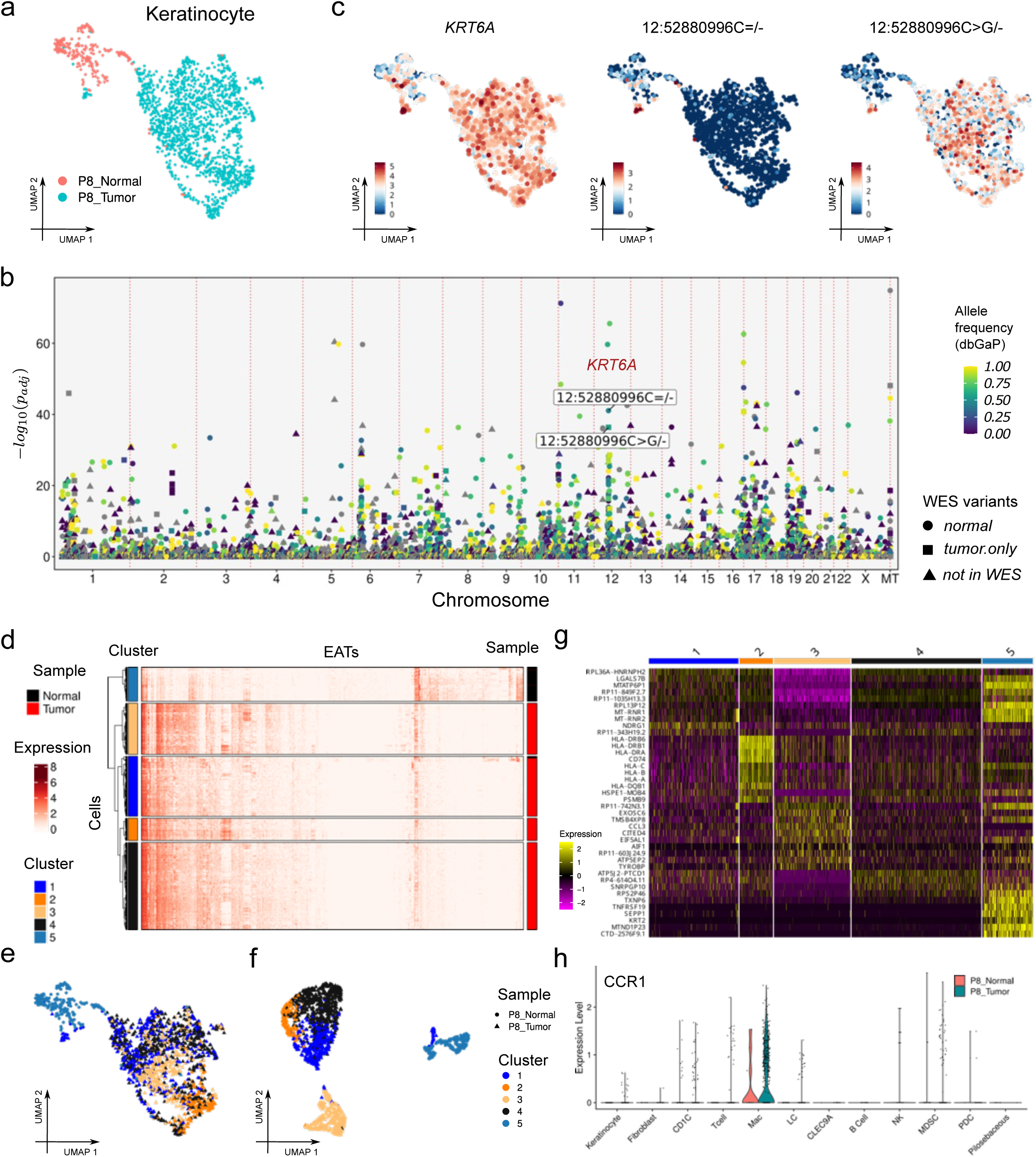
| Identification of cSCCs subclones with alternative expressed EATs. (a) UMAP visualization of keratinocytes, colored by tissue type. (b) Manhattan plot illustrating the results of the spatial dissimilarity test between EATs and other EATs at the same locus. Points are color-coded by AF from the dbGaP/ALFA database, shaped by variant origin, and sorted by genomic coordinates along the x-axis. P values for each event were transformed using a negative log10 scale for visualization. (c) UMAP plots showing the expression of gene KRT6A (left), EAT 12:52880996 allele C (middle) and allele G (right). Expression levels were retrieved from ‘data’ layer of Seurat object with default method. (d) Heatmap representing the expression of prioritized EATs across all keratinocytes. Rows correspond to cells, clustered into five groups, while columns represent EATs. (e) UMAP visualization of keratinocytes, color-coded by the cell clusters defined in the heatmap (d). Points are additionally shaped according to tissue type. (f) Refined UMAP plot of keratinocytes, further highlighting clustering based on selected EAT expression. Points are color-coded by cell clusters and shaped according to tissue type. (g) Heatmap of scaled gene marker expression across the defined cell clusters in (d). (h) Violin plot showing *CCR1* gene expression across various cell types and tissues, separated by tissue type to highlight differences in expression patterns.

Performing the SD test across all EATs (Mode 2) identified 265 EATs across 195 loci (Fig. 5b). Notably, we observed a concentration of alternatively expressed EATs at chromosome 12q13.13, predominantly originating from the *KRT16A* gene. Some EATs effectively distinguished tumor from normal cells; for example, EAT 12:52880996 allele G was expressed across all cell groups, whereas allele A was highly expressed in normal cells but sporadically in tumor cells, suggesting its utility for identifying subpopulations (Fig. 5c).

Hierarchical clustering of keratinocytes using EATs from the 195 loci resulted in five distinct groups, with most normal tissue cells grouped into cluster 5 (Fig. 5d).

Mapping these clusters onto the original UMAP did not clearly separate groups 1, 2, and 3 (Fig. 5e). However, re-clustering the cells based on alternative expressed EATs achieved clear segregation of these groups (Fig. 5f).

Further gene-level expression analysis among the clusters revealed differential expression of cancer-related genes, such as *DDIT4*^38^ and *HILPDA*^39^ across tumor subclones (Fig. 5g). Of particular interest, group 2 exhibited high expression of the ligand gene *CCL3*, while the corresponding receptor gene *CCR1* was expressed in tumor cells (Fig. 5h). The *CCL3-CCR1* axis has been implicated in tumor progression in various cancers^40,41^, suggesting its potential role in cSCC subclone dynamics.

### Spatial dissimilarity analysis of EATs in 15 adult human tissues reveals new allele-specific expressed genes

We explored ASE across 15 organs from an adult male donor (AHCA project) using 84,363 cells^42^. Cell annotations were adapted from the original study, with subclusters consolidated into major cell types (e.g., grouping all T cell subtypes as “T cells”) (Fig. 6a). From 176,883 loci, we identified 219,671 EATs, of which 2,894 showed significant alternative expression (adjusted P < 1e-10) (Fig. 6c).

**Figure 6.**
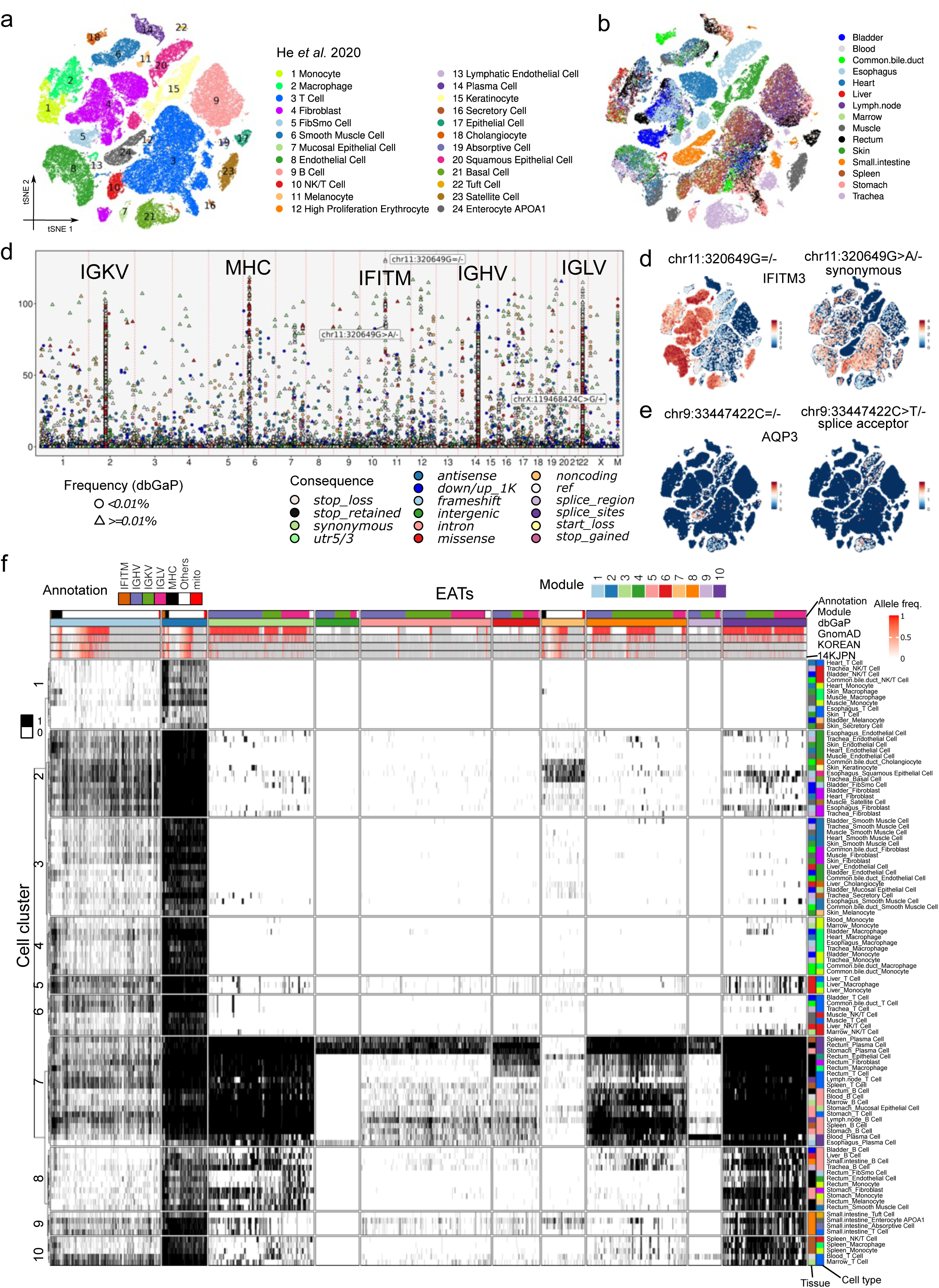
| Spatial dissimilarity analysis of EATs in 15 adult human tissues reveal new allele-specific expressed genes. (a) tSNE visualization of cells from the AHCA project, color-coded to indicate major cell types. (b) tSNE plot with cells color-coded based on their tissue or organ of origin. (c) Manhattan plot illustrating the results of the spatial dissimilarity test between EATs and other EATs at the same locus. Points are color-coded by molecular consequences and shaped by AF from the dbGaP/ALFA database and sorted by genomic coordinates along the x-axis. P values for each event were transformed using a negative log10 scale for visualization. (d) tSNE visualization of the normalized expression level of EAT chr11:320649/-allele G (left) and allele A (right). (e) tSNE visualization of the normalized expression level of EAT chr9:33447422/-allele C (left) and allele T (right). (f) Heatmap of alternatively expressed EATs, with rows clustered into 10 clusters based on merged cells and columns clustered into 10 modules based on EATs. Expression values of EATs were merged by cell type and tissue, then binarized before clustering. Within each module, EATs are sorted by AF from dbGaP/ALFA databases, and within each cluster, cell groups are sorted by type. GnomAD, the Genome Aggregation database; KOREAN, Korean Reference Genome Project; 14KJPN, Japanese population reference panel.

The Manhattan plot revealed six prominent peaks, including loci on autosomal chromosomes and mitochondria, with four peaks mapped to Ig (IGHV, IGKV, IGLV) and MHC regions (Fig. 6c, Supplementary Fig. 8a). Notably, a peak at chromosome 11p15.5 corresponded to IFITM genes (Supplementary Fig. 8b), known for their antiviral role^43^. ASE events at these loci were cell-type specific; for instance, EAT chr11:230649 allele G was highly expressed in fibroblasts and immune cells, whereas allele A showed enriched expression in T cells and varied between monocytes and macrophages (Fig. 6d).

To systematically evaluate ASE, we classified EATs into two categories: (1) Class I: Both alleles at the locus showed significant SD test results, indicating bona fide ASE with distinct expression in different cell populations. (2) Class II: Only one allele exhibited significant expression, often indicating sparse expression of the complementary allele, potentially due to somatic mutations or stochastic expression^13,14,44,45^. For example, EAT chrX:119468424G/+ was found across tissues, while its complementary allele C showed sparse expression, suggesting a somatic mutation early in embryogenesis (Supplementary Fig. 8c).

Among 5,726 significant EATs, 1,023 from 508 loci were classified as Class I, with seven loci containing >2 EATs. Most Class I EATs mapped to MHC/Ig regions, while 16 EATs were derived from IFITM genes. Additionally, 62 EATs across 24 genes showed cell-type-specific expression (Supplementary Fig. 9, 10a). Notably, these genes exhibited allele-biased expression but were not associated with known mechanisms such as imprinting, X inactivation, or somatic rearrangements. For example, the macrophage marker gene *CD68* displayed promoter-specific ASE in fibroblasts^46^, where EAT chr17:7579637 allele T was enriched, while allele G showed distinct patterns in myeloid cells (Supplementary Fig. 10a).

EAT consequence predictions revealed instances of functional impact. For instance, chr9:33447422 allele T in the splice acceptor region of *AQP3* was absent in enterocytes, suggesting elimination of mutated cells. This aligns with impaired enterocyte proliferation observed in *AQP3* deficiency^47^. Additionally, we identified mutations altering protein structure expressed in cell subclones (Supplementary Fig. 10b, c).

To assess ASE distribution, we aggregated cells into pseudo-bulk samples by cell types and tissues (_≥_50 cells/sample). Clustering analysis of 5,420 EATs identified 10 clusters of samples and 10 EAT modules (Fig. 6f). Ig variants were enriched in modules 1, 2, and 7, while modules 3 and 10 contained EATs with high AF in the dbGaP/ALFA and other databases, suggesting reference allele expression. Interestingly, fibroblasts from the rectum and stomach expressed module 2 and 10 EATs, differing from fibroblasts in the heart, skin, and muscle, suggesting organ-specific ASE patterns.

This analysis highlights the value of ASE in understanding gene regulation across tissues and cell types. Novel ASE genes, influenced by promoter usage and variant consequences, were identified, expanding our understanding of ASE mechanisms. Combined with AF databases, our approach also uncovers informative somatic mutations, offering deeper insights into gene function and lineage development in normal human tissues.

## Discussion

In this study, we extended spatial statistics to single-cell analysis by utilizing a cell-cell weight matrix and addressing the spatial dissimilarity challenge with a novel spatial measure. This method examines distinct spatial expression patterns between two features in single cells, focusing solely on the spatial autocorrelation of the test feature while leaving the binding feature unexamined. Statistical significance is assessed via a permutation test, which permutes only the test feature’s distribution, ensuring robust evaluation.

The D score integrates two components: L, which measures the spatial autocorrelation of the test feature, addressing scRNA-seq sparsity by evaluating spatial distribution instead of expression levels, and 1 - r which quantifies dissimilarity between the test and binding features, overcoming challenges such as low gene body coverage. Features with low expression but spatial autocorrelation and distinct expression patterns relative to their binding features can be prioritized, making the method particularly effective for biased scRNA-seq datasets, the most widely used single-cell sequencing technology.

Our method requires only feature counts and a cell-cell weight matrix, eliminating the need for prior cell clustering - a commonly non-robust step due to different resolution in cell cluster definitions^15,16^. The flexibility in defining binding relationships makes the SD test applicable to diverse studies beyond AS and ASE, such as analyzing spliced versus unspliced gene expressions to study gene expression dynamics. Another innovative feature of our SD test is its integration with Manhattan plots, akin to GWAS, enabling the identification of genomic hotspot regions with significant alternative expression differences. The SD test also supports ASE analysis and cell lineage tracing using expressed EATs, which act as natural barcodes for tracking developmental trajectories in human normal and cancer tissues without prior read assembly. Yano, our R package, further includes tools for genetic variant annotation using various databases and functional predictions.

By applying the SD test and genetic variants annotation to single-cell data from the same tissue types across individuals, we can explore gene functions at specific cell types. With the upcoming release of the first draft of the Human Cell Atlas^48^, we anticipate that future research will focus on advancing gene expression analysis to the isoform level and linking genotype to phenotype through population-based cell atlases. Our method and tools are poised to address these emerging challenges.

## Data availability

Alignment BAM files and FASTQ files for MOp datasets were downloaded from NeMO Archive (https://assets.nemoarchive.org/dat-ch1nqb7). Raw published data for cSCC and adult human datasets are available from GEO under accession no. GSE144240 and GSE159929 respectively. Mouse reference database and annotation were obtained from the 10X Genomics resource page with the code refdata-gex-mm10-2020-A. Human reference database and annotation were downloaded from the 10X Genomics resource page, using the code refdata-cellranger-hg19-3.0.0 for the hg19 human reference and refdata-gex-GRCh38-2020-A for the GRCh38 reference. NCBI RefSeq annotation file was downloaded from https://hgdownload.soe.ucsc.edu/goldenPath/archive/hg38/ncbiRefSeq/110/hg38.110. ncbiRefSeq.gtf.gz and was used to predict molecular consequence of genetic variants. Population based AF database was downloaded from https://ftp.ncbi.nih.gov/snp/latest_release/VCF/GCF_000001405.40.gz for GRCh38 and https://ftp.ncbi.nih.gov/snp/latest_release/VCF/GCF_000001405.25.gz for hg19, and the chromosome names and type of tags were manually converted. The ClinVar database was downloaded from https://ftp.ncbi.nlm.nih.gov/pub/clinvar/vcf_GRCh38/archive_2.0/2024/clinvar_2024 0317.vcf.gz for GRCh38 and https://ftp.ncbi.nlm.nih.gov/pub/clinvar/vcf_GRCh37/archive_2.0/2024/clinvar_2024 0312.vcf.gz for hg19.

## Code availability

The package is open-source and available at https://github.com/shiquan/Yano. Codes for this paper can be reached at https://github.com/shiquan/Yano_paper.

## Acknowledgments

We thank Albin Sandelin, Rickard Sandberg, and Lin Lin for discussions and comments on the early versions of this work.

## Contributions

Q.S. and K.K. conceived the project. Q.S. developed the concept, implemented the code, performed the analyses, and drafted the manuscript. K.K. supervised the work and reviewed and edited the manuscript.

## Methods

### Feature annotation and counting with PISA

The feature annotation process begins with BAM files, using the enhanced functionality of the PISA^26^ software. New parameters, -exon and -psi, have been added to the PISA anno (v1.2). By default, the annotation process is restricted to reads that are entirely enclosed within exonic regions. Gene names are recorded under the GN tag, while exon names are documented under the EX tag. For reads spanning junctions or multiple exons, an additional JC tag is generated. Excluded exons are represented with the ER tag. The naming convention for exons, junctions, and excluded exons is consistent and follows the format: chromosome:start-end/strand/gene. In cases where strand specificity is not applicable, such as non-strand-sensitive libraries, the strand field is omitted. Notably, exon and exon-excluded reads share the same name but have distinct meanings: (1) Exon counts represent reads entirely enclosed within exons or junction reads overlapping the exon; (2) Exon-excluded counts represent junction reads that exclude the exon.

For SNV annotation, the -ref-alt parameter has been introduced in PISA anno (v1.2). By default, the annotation process is strand-sensitive; however, for non-strand-sensitive data (e.g., Smart-seq2), the -is parameter (short for “ignore strand”) can be used. The name for EAT follows the format: “chromosome:pos/reference allele > alternative allele/strand” for SNVs or “chromosome:pos/reference allele=/strand” for reference allele, the strand field is omitted when -is set.

After feature annotation, the PISA count function quantifies UMI counts for each feature within individual cells. For example, gene expression counts can be obtained using the following command:

“PISA count -tag CB -umi UB -outdir exp -anno-tag GN anno.bam”.

In this command, CB specifies the cell barcode; UB specifies the UMI; -anno-tag determines the feature type to count, such as genes (GN), exons (EX), junctions (JC), excluded exons (ER), or EATs (VR).

### Implementation of the spatial dissimilarity test method

Our approach comprises a multi-step process to perform the spatial dissimilarity test of features within single cell space.

Step 1: Preparing the Seurat object. A Seurat object is prepared using gene expression counts. Following standard preprocessing steps, including normalization, selection of HVGs, scaling of HVGs, and PCA analysis. Cell clustering is optional for this analysis but can usually be performed for visualization. In this workflow, the GetWeights function in the Yano package is utilized to create a shared nearest neighbor (SNN) graph using Seurat’s FindNeighbors function. This SNN graph is then normalized column-wise, ensuring that the sum of each column is one, thereby forming a weight matrix graph. This normalized graph is essential for subsequent calculations, serving as the foundation for evaluating the spatial autocorrelation of each feature in the next step.

Step 2: Incorporating additional features into Seurat object. Next, we incorporate other features (e.g., exons or junctions) as separate assays within the Seurat object using CreateAssayObject. We then switch the default assay to the test assay. By executing RunAutoCorr from Yano, Moran’s I is calculated for the test features. The Moran’s I is defined as

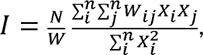

where *N* is the cell number, and *W* represents the sum of all values in the weight matrix. Consider that the sum of each row is always equal to 1, and 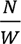 always should be 1, then, the formula simplifies to

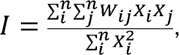

with P values derived from the Z scores calculated by the standard deviation of I^30^. P values are then adjusted with the Benjamini-Hochberg procedure^31^. Features with significant spatial autocorrelation are identified using function SetAutoCorrFeatures in Yano. The default cutoff for spatial autocorrelated features is Moran’s I > 0 and adjusted P value < 0.01.

Step 3: Feature annotation and spatial dissimilarity testing. Specific functions from Yano, like ParseExonName and ParseVarName, are designed to parse the genomic coordinates from the feature names which generated with PISA. The RunBlockCorr function conducts spatial dissimilarity tests on selected features with binding assay. This step involves setting binding relationships between features and calculating the D score, which combines spatial smoothing and dissimilarity measures:

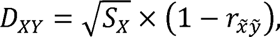

where *X* is the test feature, *Y* is the binding feature. *S_X_* is a spatial smooth scalar of test feature, defined as

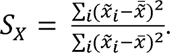

*S_X_* is used to describe spatial association degree^21^. *X̃* is the spatial lag value of *X* by weight matrix *W*. The 1-Pearson correlation coefficient *r* describe the dissimilarity between spatial lag of *X* and *Y*. Therefore, the *D* score integrates the spatial smoothing degree of test feature and the dissimilarity between test feature and the binding feature. Yano then permutates the order of *x* and calculate the *D* score 100 times to evaluate the expected value and variance, and estimate significance, with P values calculated by one-tail *t*-test via the ‘pt’ function from the R stats package. P values were automatically adjusted using the Benjamini-Hochberg procedure, implemented with the ‘p.adjust’ function from the R stats package. Finally, the FbtPlot function visualizes these P values on a Manhattan plot.

### Visualization of single cell alignments with track plots

To elucidate the distribution of single-cell alignments across genomic regions, including junction reads, we have implemented a track plot visualization tool, TrackPlot, within Yano. TrackPlot provides an intuitive representation of read coverage across different cell groups, leveraging UMI depth as a metric. The process begins with the preparation of a sorted and indexed BAM file. The track plot function then computes the UMI depth at each genomic coordinate. To enhance visualization performance and achieve a smoother appearance of the tracks, the target region is segmented into 1,000 bins. The average UMI depth across these bins is calculated and utilized for the plot.

To account for variations in group sizes, the UMI depth for each cell group is normalized to the number of cells in the group, resulting in a normalized UMI depth per cell. This step ensures that the tracks are directly comparable between groups, allowing for a clear visual comparison of alignment distributions. But if cell groups are not specified, the track plot will generate raw UMI depth for all cells.

By default, TrackPlot distinguishes the strand of reads during visualization. UMI depth for the reverse strand is adjusted by applying a negative conversion, and reads are color-coded by strand, with forward strand reads displayed in red and reverse strand reads in blue. When the UMI tag is not set, TrackPlot defaults to using raw read depth instead of UMI depth for visualization. If the strand parameter is set to ‘ignore’, reads from both strands are summed up, providing a combined view without strand differentiation.

For analyses involving multiple sequencing runs or datasets, Yano supports the input of a list of BAM files, allowing for the concurrent visualization of alignment tracks from several samples or conditions. This feature is particularly useful for comparing and contrasting alignment patterns across different sequence libraries, samples or experimental groups.

### EAT calling

For the sorted and indexed BAM files, bcftools is employed for genetic variant calling. Given the intrinsic non-strand-specific nature of DNA variant callers, the standard filtering methods typically used for DNA may not be suitable for RNA datasets due to potential strand biases, particularly in the context of ASE. To address this, the unfiltered, raw variants are used during the EAT annotation phase. The annotation of all detected alleles at a given locus with -ref-alt parameter. Additionally, PISA anno enables the strand-sensitive annotation of EATs in default. For paired-end sequencing data, the strand information for EATs from the second read is inverted to accurately reflect the original cDNA fragment’s strand.

### EAT annotation

The annoVAR function in the Yano package is designed to annotate EATs using various databases and predict their molecular consequences based on gene annotation files. This function accepts essential inputs such as EAT locations, reference and alternative alleles, and strand information. When using databases in VCF format, the tags parameter must be specified to extract desired fields, adhering strictly to VCF specifications (VCFv4.2, https://samtools.github.io/hts-specs/VCFv4.2.pdf). For instance, records with a ‘Number’ type of A, which stores allele-specific information separated by commas, the function extracts data exclusively for the alternative allele.

For molecular consequence predictions, the function requires a GTF annotation file to define gene and transcript boundaries and an indexed reference genome in FASTA format. Unlike DNA variant annotations, annoVAR accounts for strand-sensitive RNA reads, as different transcripts or genes on opposite strands can produce distinct consequences for the same allele. The function predicts a consequence for each overlapping transcript and ranks them by impact, exporting only the most significant annotation and gene name for simplicity in downstream analysis. The ranking prioritizes coding genes over noncoding and antisense genes for reference alleles, while for non-reference alleles, a detailed ranking system places highly impactful consequences. Consequences are ranked in the following order: whole_gene > exon_loss > stop_gained > exon_splice_sites > splice_donor > plice_acceptor > frameshift_truncation > frameshift_elongation > stop_loss > start_loss > inframe_indel > missense > synonymous > stop_retained > start_retained > utr5 > utr3 > splice_region > intron > utr5_intron > utr3_intron > noncoding_exon > noncoding_splice_region > noncoding_intron > upstream_1K > downstream_1K > antisense_utr3 > antisense_utr5 > antisense_exon > antisense_intron > antisense_upstream_1K > antisense_downstream_1K > intergenic.

Moreover, if a chromosome is missing in the annotation file, the consequence is marked as ‘unknown’. For non-strand-sensitive EATs, the antisense records will not be generated.

### Analysis of mouse primary motor neuron data

The raw FASTQ files for the Smart-seq2 data were downloaded from NeMO Archive. Reads for each cell were aligned to the mm10 reference genome using STAR (v2.7.1a), with the RG tag added during alignment. The resulting BAM files were sorted and merged using sambamba (v0.8.0). The sorted and indexed BAM files were subsequently used for feature annotation and counting. Feature annotation was performed with PISA (v1.2), using the -exon, -psi, and -is parameters. A GTF file (gencode vM23) downloaded from the 10X Genomics website was used to annotate genes and exons. Non-strand-sensitive features were counted with PISA count, with the -cb parameter set to RG. No UMI tag was used for Smart-seq2 data.

The resulting gene count file was loaded into the R environment using the ReadPISA function from the Yano package. The loaded count matrix was converted into a Seurat object and normalized using a log-scaled approach, followed by adjustment for library size, as implemented in the Seurat (v5.1.0) pipeline. Default parameters and the top 20 PCs were used for cell clustering and dimensional reduction analysis.

Counts for exons, junctions, and exon-excluded reads were loaded into the Seurat object as new assays using the CreateAssayObject function from the Seurat package. Features detected in fewer than 20 cells were excluded. Raw counts were normalized using Seurat’s NormalizeData function. The top 20 PCs calculated with the gene assay were used to construct a cell-cell similarity matrix, which was subsequently employed to calculate Moran’s *I* and perform spatial dissimilarity analysis follow procedures of “Implementation of spatial dissimilarity test method”.

For alternative expression analysis with exon and junction assays, Mode 1 is used for comparison with gene assay. While Mode 3 is used when comparison between exon assay and exon excluded assay.

As most published scRNA-seq datasets are generated with strand-sensitive kits, both PISA and Yano enable strand-sensitive processing by default. Track plot visualization with the TrackPlot function by setting the UB parameter to NULL, CB parameter set to RG and the strand parameter was set to ‘ignored’ to accommodate Smart-seq2 data.

The aligned BAM files for the 10X Genomics v3 data were directly downloaded from NeMO Archive. Feature annotation was performed with PISA (v1.2), using the -exon and -psi parameters. During feature counting, the CB tag was used to identify cell barcodes, and the UB tag was used to count UMIs. Post-processing for the 10X data was carried out using the same Seurat and Yano workflow as described for Smart-seq2 data. During visualization of track plots for 10X data, default parameters were used.

### Cluster based alternative splicing analysis with DEXSeq and BRIE2

To run DEXSeq on selected cells, we implemented the RunDEXSeq function in the Yano package. The process involves generating pseudo-bulk groups from single-cell raw feature counts for each cell group. For each comparison, three pseudo-bulk samples are generated for each group by aggregating raw feature counts from the single cells. The group 1, specified with ‘ident.1’, is the primary cell group for comparison. By default, all cells not in group 1 are aggregated into group 2 unless explicitly specified with ‘ident.2’. A count matrix of the test assay with six pseudo-samples (three for each group) is prepared as the input for DEXSeq. Simultaneously, for the binding assay, six pseudo-samples from the same cells are generated from the raw counts of the binding assay and used as alternativeCountData in DEXSeq.

DEXSeq is run using a single thread, which makes the process slower when analyzing all features. To optimize runtime, only spatially autocorrelated features are input for benchmarking purposes.

For the BRIE2 pipeline, the PISA pick module is used to split the merged BAM file into a one-cell-per-file format, as this is the required input format for brie-count. The brie-count and brie-quant pipelines were executed according to the standard protocols provided in the BRIE2 manual (https://brie.readthedocs.io/).

### Analysis of squamous cell carcinoma data

Aligned BAM files were downloaded from Gene Expression Omnibus (GEO) datasets, adopting filtered cells and annotations directly from the original studies. Due to discrepancies in the versions of the human reference and mitochondrial genomes used in WES and scRNA-seq datasets, WES reads were realigned to the hg19 human reference genome using BWA. This was followed by the repair of paired-end information and marking of duplicate reads using samtools.

Genetic variants in both tumor and normal libraries were identified using bcftools. This process resulted in four variant files in VCF format, two each from scRNA-seq and WES datasets. These files were merged, and indels were normalized using bcftools. The merged and normalized VCF file was used to annotate the BAM files employing PISA.

The quantified gene expression data served as the basis for cell clustering, executed via the Seurat package employing its default parameters. The EAT counts read with ReadPISA and load into Seurat as a new assay. Features detected in fewer than 10 cells were excluded from downstream analysis to maintain data quality and relevance. We then performed spatial dissimilarity analysis follow procedures of “Implementation of spatial dissimilarity test method”.

### Analysis of adult human cell atlas

Raw FASTQ files were downloaded and processed using the CellRanger pipeline (version 6.1.2) with ‘SC5P-PE’ chemistry. This step produced BAM files, which were then utilized for subsequent annotation and feature counting phases. To maintain consistency with previous findings and leverage established insights, filtered cells, cell annotations, and tSNE maps were directly adopted from the original study.

Genetic variants for each tissue sample following procedure of section “EAT calling”. To ensure the robustness of downstream analysis, features detected in fewer than 20 cells were filtered out. The gene expression data was employed for PCA analysis using the Seurat package, adhering to its default settings.

EAT counts were read and loaded as a new assay for Seurat object. The top 20 PCs built by gene expression were then used to build cell-cell weight matrix. The locus names and positions of EATs were parsed with ParseVarName function. The UCSC RefSeq gene annotation for GRCh38 was used to predict the consequences of EATs with annoVAR.

## Supplementary Figure Legends

**Supplementary Figure 1.**
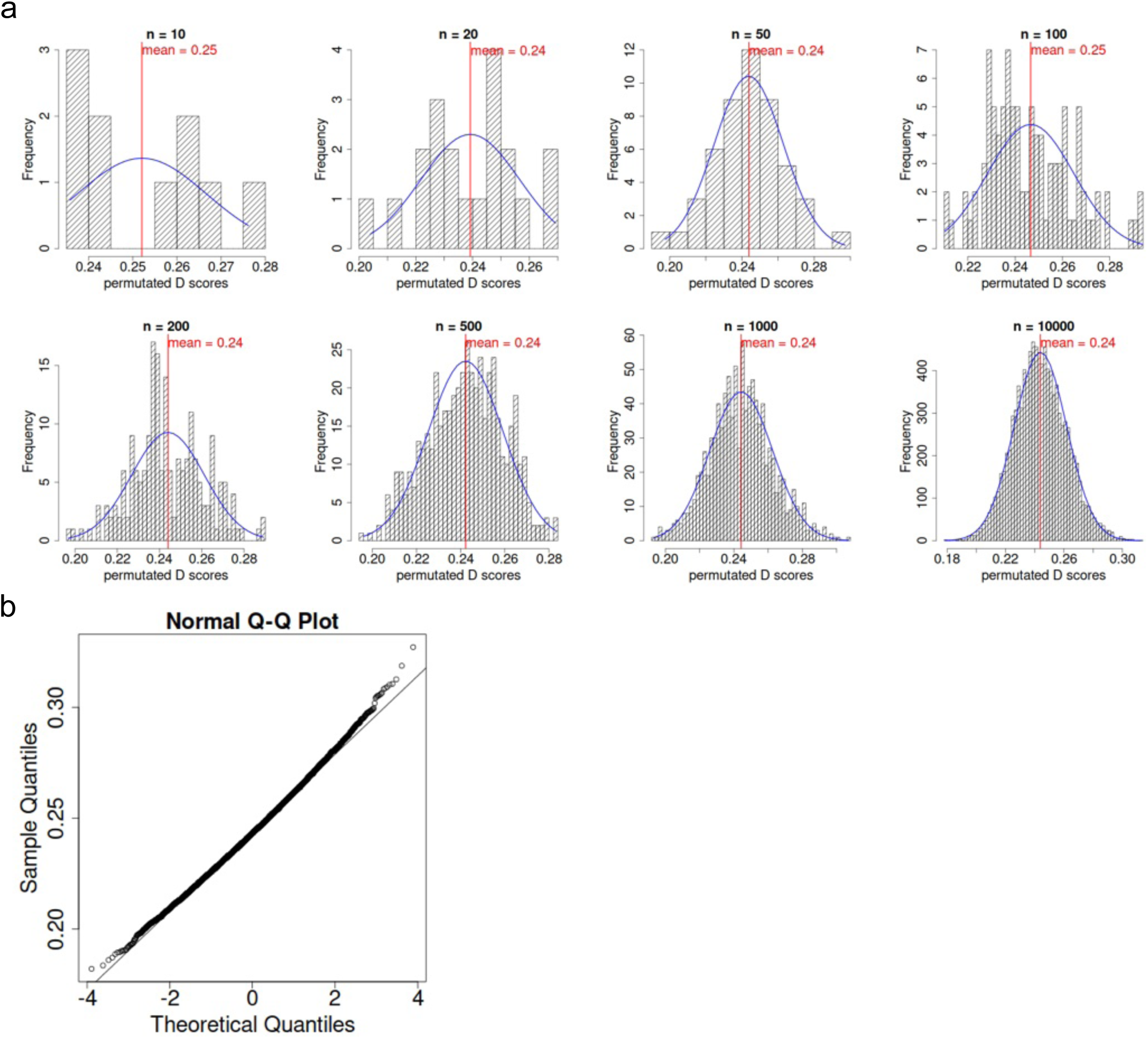
| Analysis of D score distribution. Related to Figure 1. (a) Histogram plots showing the distribution of D scores obtained through permutation testing, ranging from 10 to 10,000 iterations. Overlaid on the histograms are curves representing the fitted normal distribution, providing a visual comparison between the empirical data and the theoretical model. This analysis validates the assumption that permuted D scores follow a normal distribution, supporting the accurate interpretation of spatial dissimilarity tests. (b) Quantile-Quantile (QQ) plot of D scores derived from 10,000 permutations, comparing the observed distribution to the expected normal distribution.

**Supplementary Figure 2.**
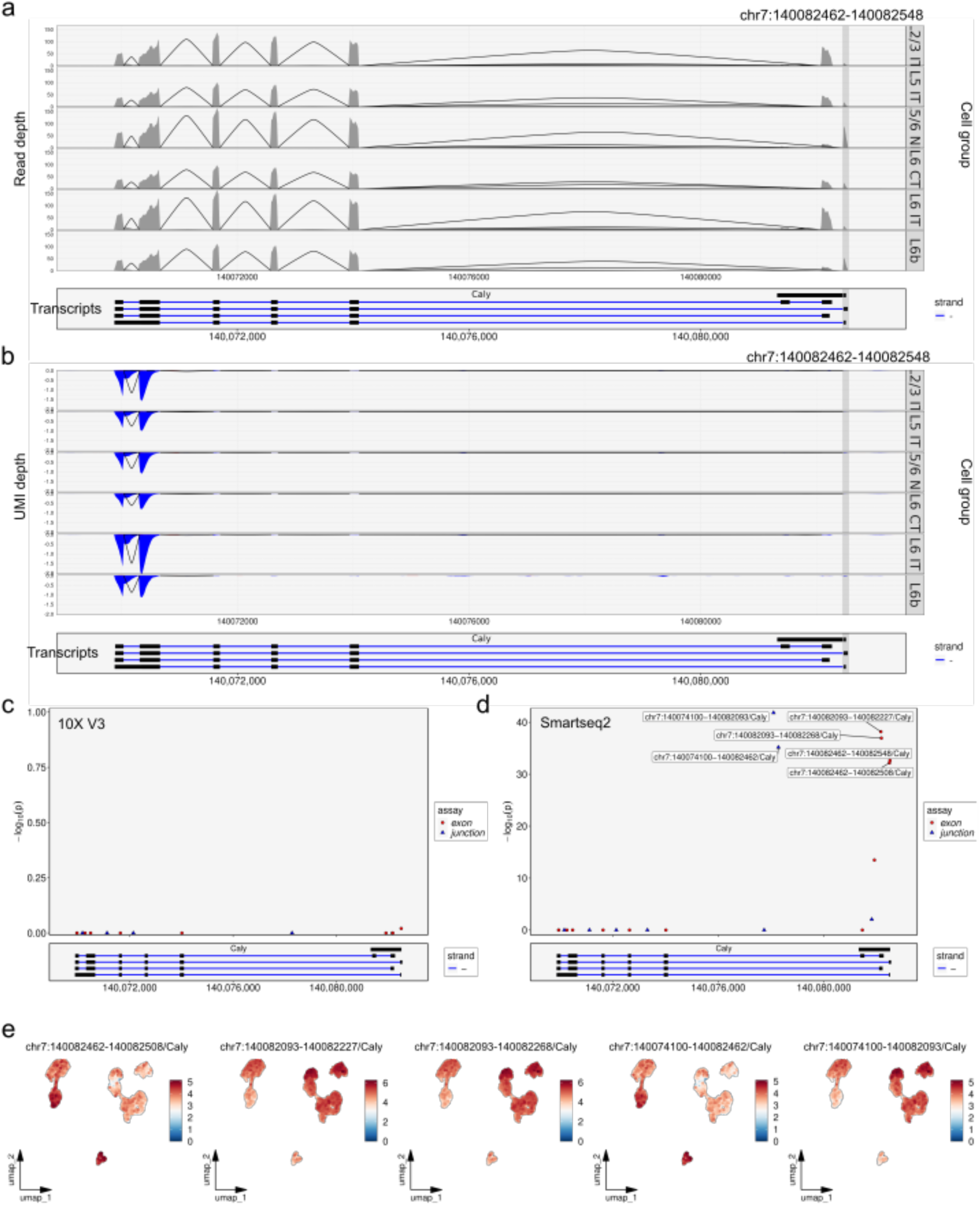
| Expression of exons and junctions of the gene *Caly* in Smart-seq2 and 10X Genomics v3 data. Related to Figure 2. (a) Track plot of normalized reads in the *Caly* gene region for Smart-seq2 data. (Top) For each cell group, read depth is divided by the number of cells, with peaks representing the mean read depth per cell within the group. Connecting lines between exons indicate the mean depth of junction reads. (Bottom) Transcript annotation for the *Caly* gene. Exon chr7:140082462-140082548 is highlighted. (b) Track plot of normalized reads in the *Caly* gene region for 10X data. (Top) For each cell group, UMI depth is divided by the number of cells, with peaks representing the mean UMI depth per cell within the group. Peaks toward the bottom indicate reads aligned to the minus strand (color in blue). Connecting lines between exons indicate the mean UMI depth of junction reads. (Bottom) Transcript annotation for the *Caly* gene. Exon chr7:140082462-140082548 is highlighted. (c) Zoom-in on gene *Caly* for Manhattan plot (Fig. 2b). (d) Zoom-in on gene *Caly* for Manhattan plot (Fig. 2c). (e) UMAP visualization of the normalized expression of exons from the *Caly* gene in Smart-seq2 data.

**Supplementary Figure 3.**
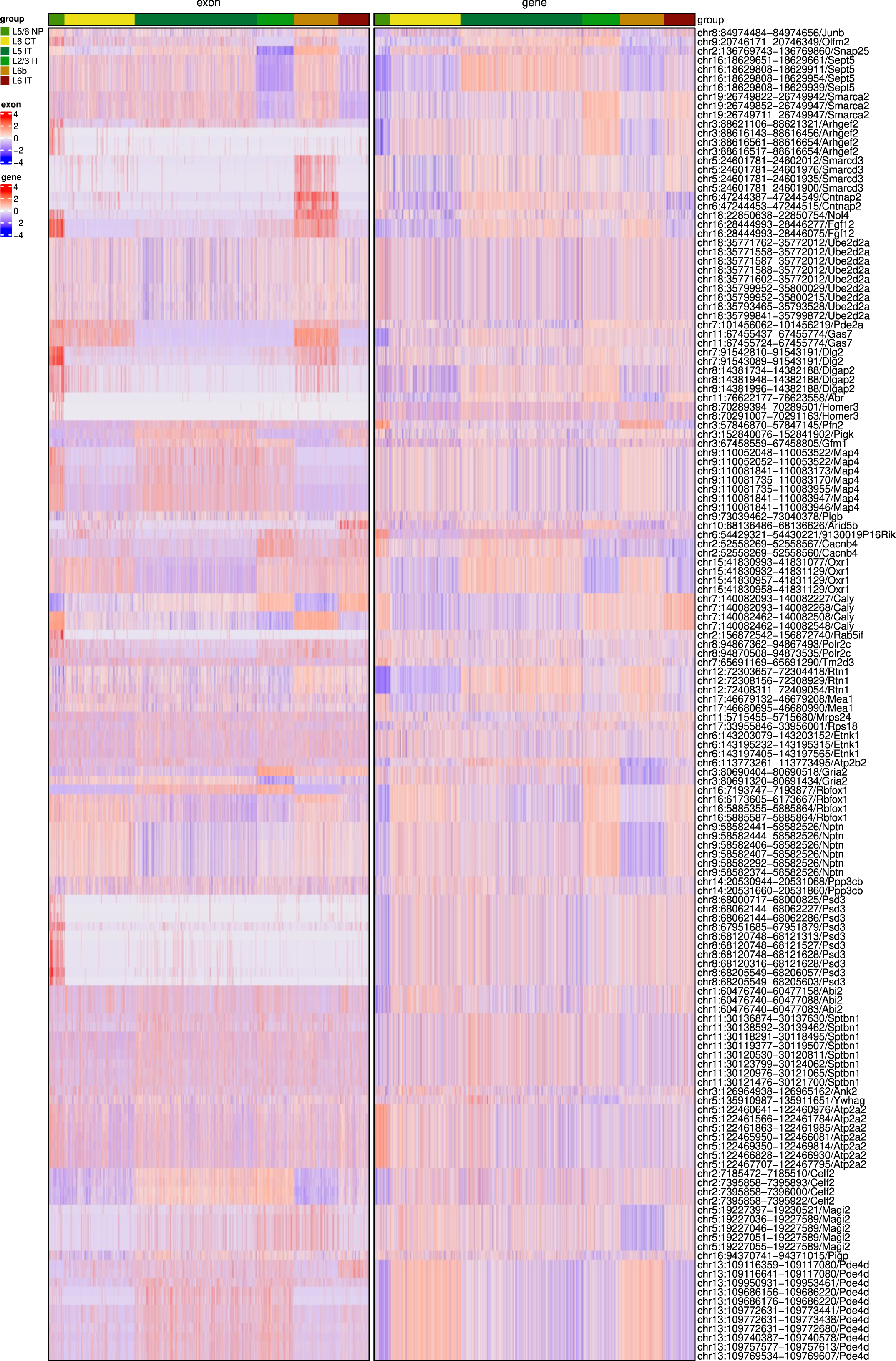
| Module analysis of alternative expressed exons for Smart-seq2 data. Related to Figure 2. Expression of exons with extremely low P values (<1e-30) are scaled and co-clustered with scaled expression of their corresponding. (Left) Heatmap of scaled exon expression. (Right) Heatmap of scaled gene expression.

**Supplementary Figure 4.**
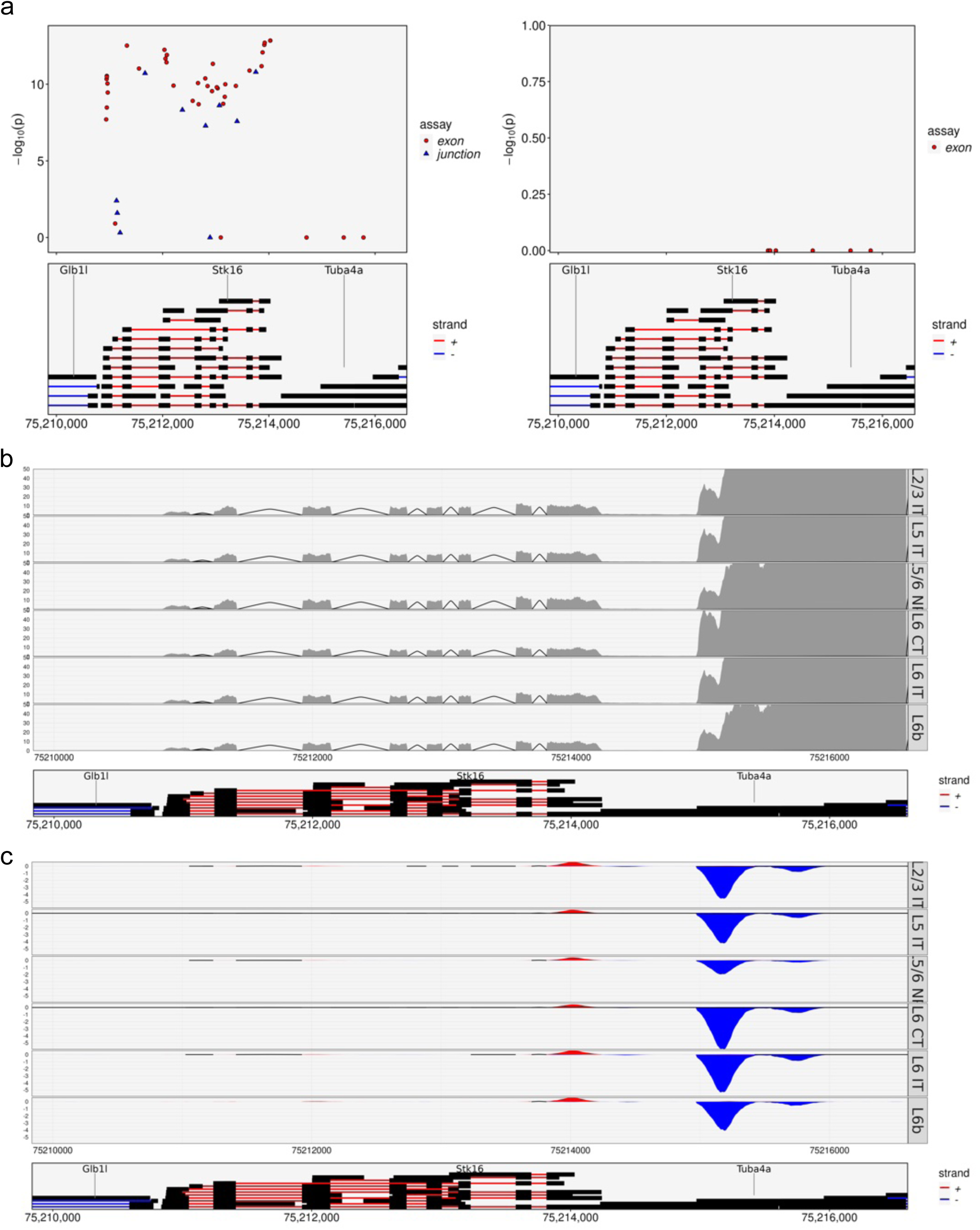
| Non-strand-sensitive data introduce bias due to overlapping genes. Related to Figure 2. (a) Zoom-in on gene *Stk16* for Manhattan plot (Fig. 2b). (b) Zoom-in on gene *Stk16* for Manhattan plot (Fig. 2c). (c) Track plot of normalized reads in the *Stk16* gene and downstream region for Smart-seq2 data. Normalized depth for each cell group capped to 50 to visualize low expression of gene *Stk16*. (d) Track plot of normalized reads in the *Stk16* gene and downstream region for 10X data.

**Supplementary Figure 5.**
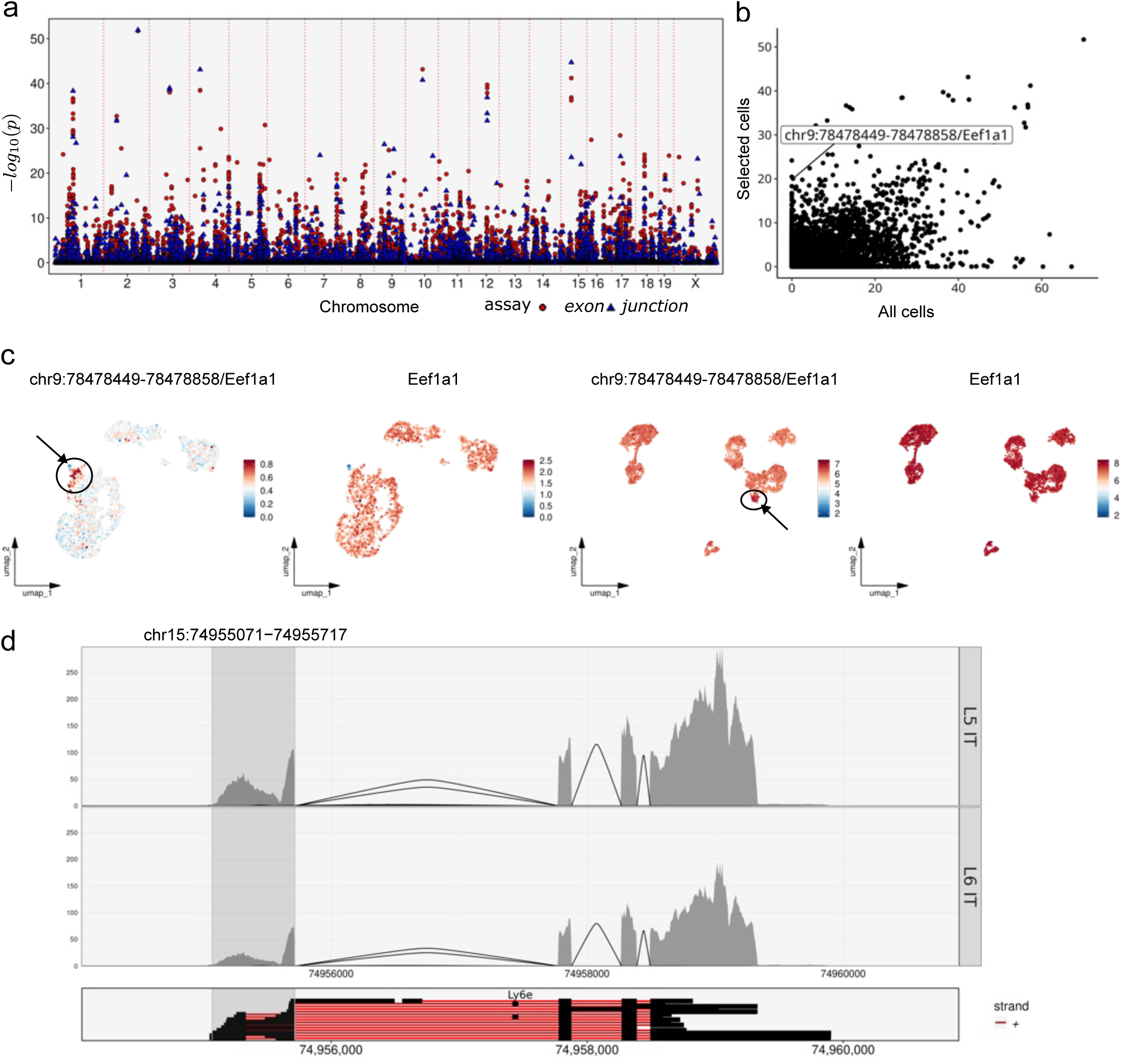
| Comparison of SD test results between selected cells and all cells. Related to Figure 3. (a) Manhattan plot illustrating the spatial dissimilarity test results between exon/junction and gene pairs for L5 IT and L6 IT cells from Smart-seq2 data. Points are color- and shape-coded based on assay type and sorted by genomic coordinates along the x-axis. P values for each event were transformed using a negative log10 scale for visualization. (b) Comparison of SD test results for the same exon analyzed with selected cells and all cells. P values are shown on a negative log10 scale. (c) UMAP plots showing the normalized expression levels of exon chr9:78478449-78478858 and its associated gene *Eef1a1* in selected cells (left two panels) and all cells (right two panels). (d) Track plot of normalized reads for the *Ly6e* gene from Smart-seq2 data. Exon chr15:74955071-74955717 is highlighted.

**Supplementary Figure 6.**
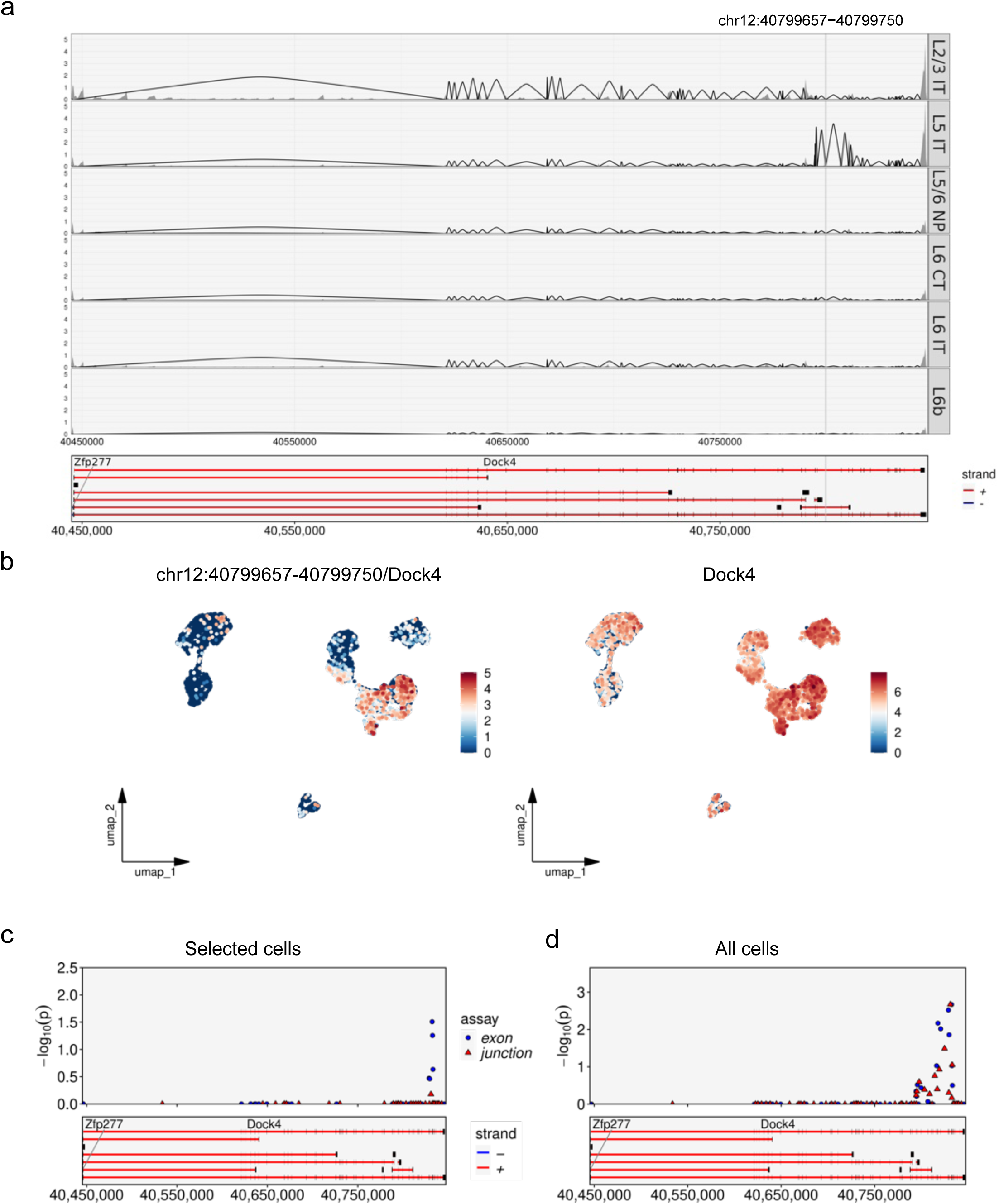
| Expression and track plot of the gene *Dock4* at all cells and selected cells. Related to Figure 3. (a) Track plot of normalized reads for the *Dock4* gene from Smart-seq2 data. Exon chr12:40799657-40799750 is highlighted. (b) UMAP plots showing the normalized expression levels of exon chr12: 40799657-40799750 (left) and its associated gene *Dock4* in all cells (right). (c) Zoom-in on gene *Dock4* region from the Manhattan plot in Supplementary Fig. 5a. (d) Zoom-in on gene *Dock4* region from the Manhattan plot in Fig. 2c.

**Supplementary Figure 7.**
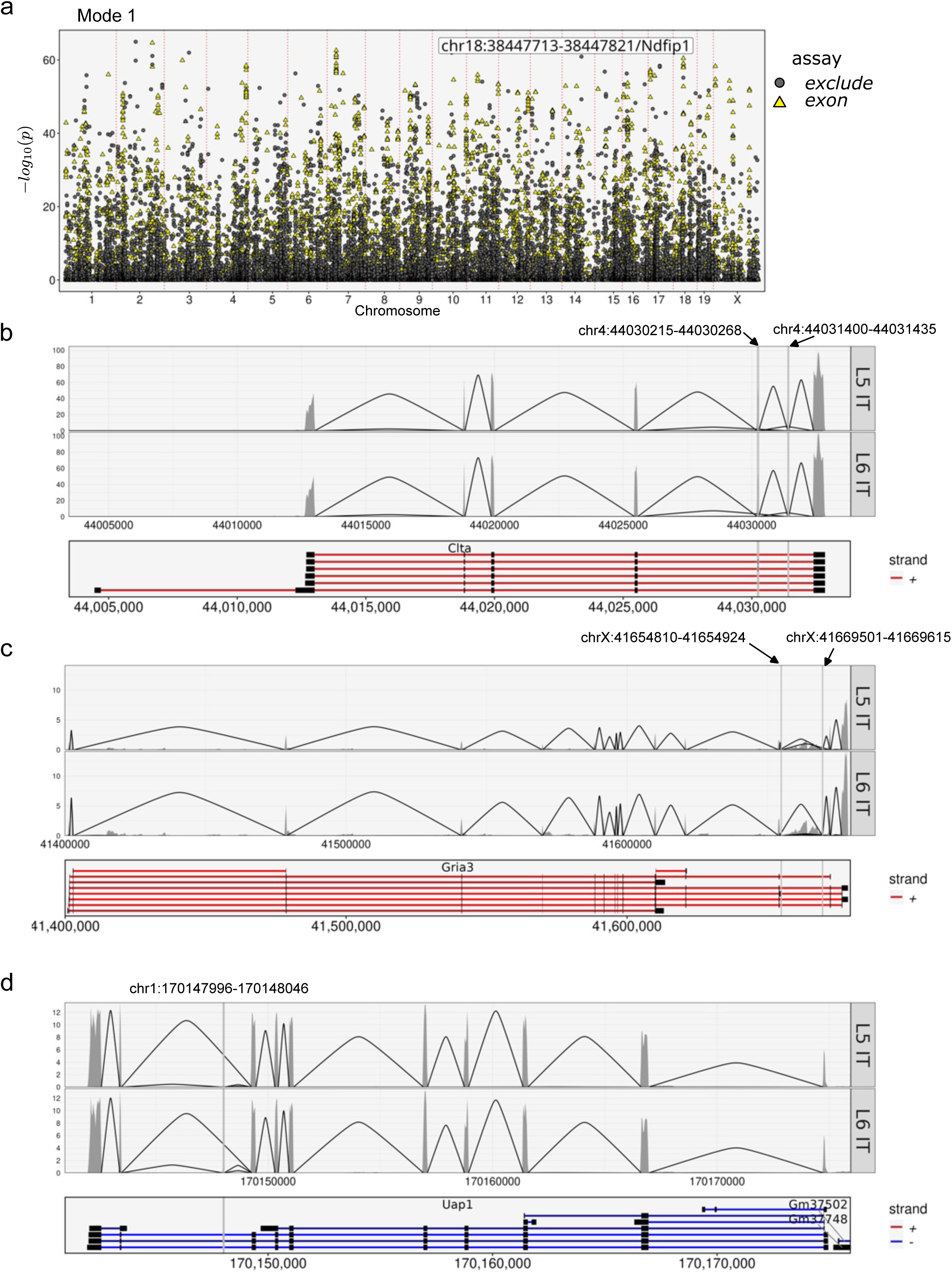
| Supplementary data for comparison with BRIE2. Related to Figure 3. (a) Manhattan plot illustrating the results of the spatial dissimilarity test between exon assay and exon-excluded assay (and vice versa) in Mode 1. Points are color- and shape-coded based on assay type and sorted by genomic coordinates along the x-axis. P values for each event were transformed using a negative log10 scale for visualization (b) Track plot showing normalized reads in the *Clta* gene region for Smart-seq2 data. For each cell group, raw depth is divided by the number of cells, with peaks representing the mean depth per cell within the group. Connecting lines between exons indicate the mean depth of junction reads. The sum of junction reads that skip an exon represents the excluded reads for that exon. Exon chr4:44030215-44030268 and exon chr4:44031400-44031435 are highlighted. (c) Track plot of normalized reads in the *Gria3* gene region for Smart-seq2 data. Exon chrX:41654810-41654924 and exon chrX:41669501-41669615 are highlighted. (d) Track plot of normalized reads in the *Uap1* gene region for Smart-seq2 data. Exon chr1:170147996-170148046 is highlighted.

**Supplementary Figure 8.**
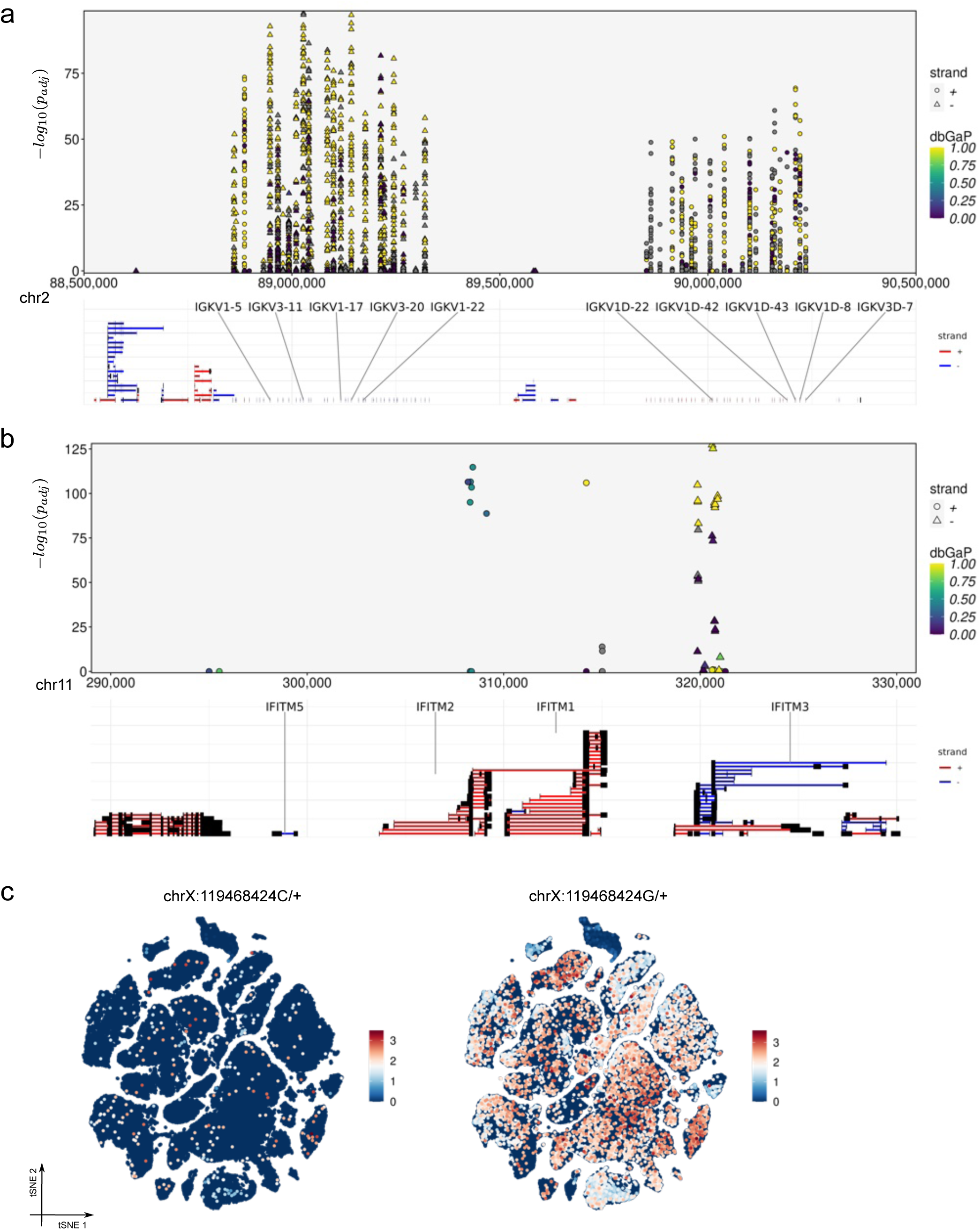
| Supplementary data supporting Figure 6. (a) Zoom-in view of the immunoglobulin kappa chain variable region from the Manhattan plot (Fig. 6c). (b) Zoom-in on IFITM genes from the Manhattan plot (Fig. 6c). (c) tSNE visualization showing the normalized expression levels of EAT chrX:119468424, with allele C (left) and allele G (right).

**Supplementary Figure 9.**
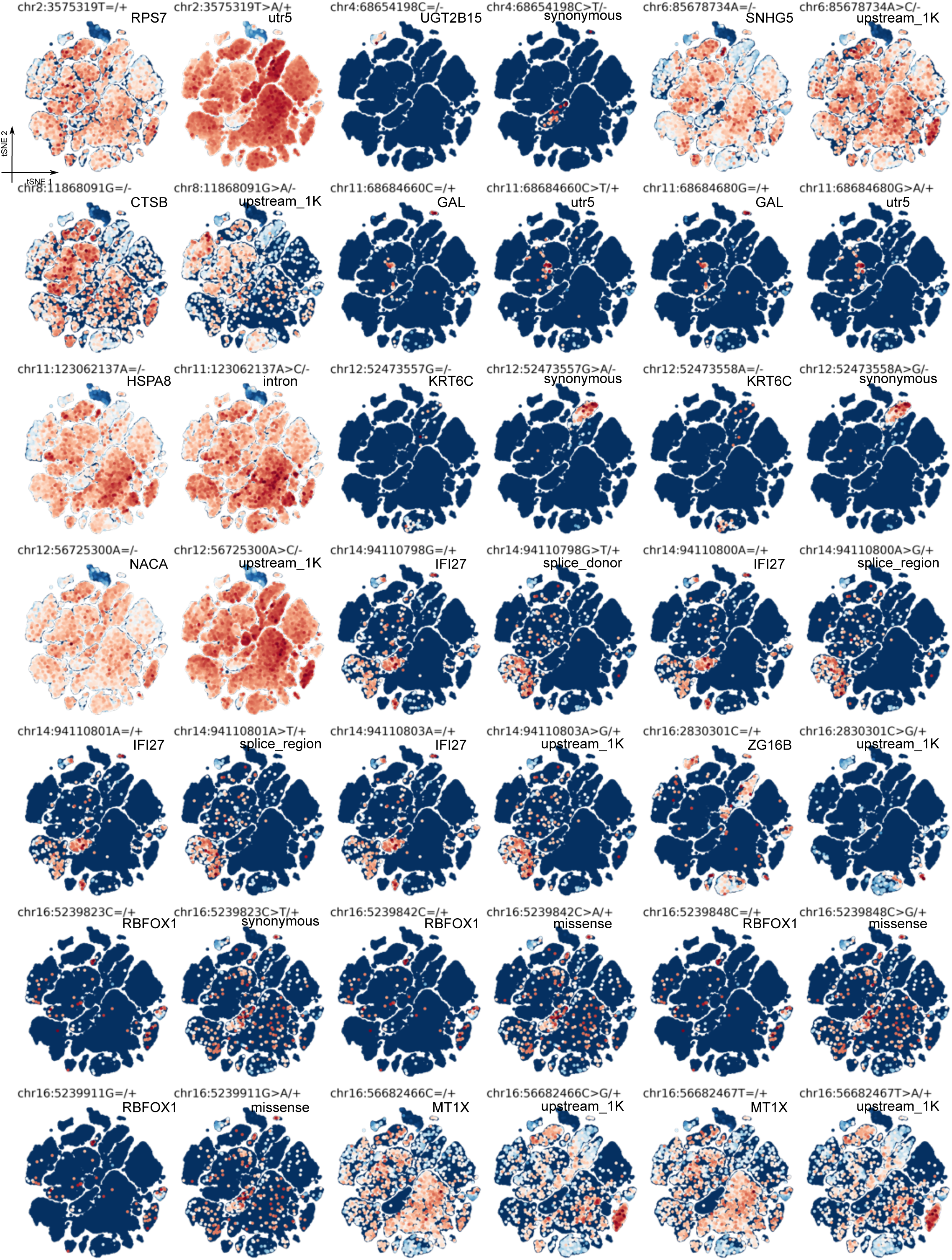
| Cases of allele-specific gene expression. Related to Figure 6. tSNE visualizations showing the normalized expression levels of EATs. Each row represents two pairs of EATs: the reference allele (left) and the alternative allele (right). The corresponding gene and molecular consequence predictions are labeled beneath the titles.

**Supplementary Figure 10.**
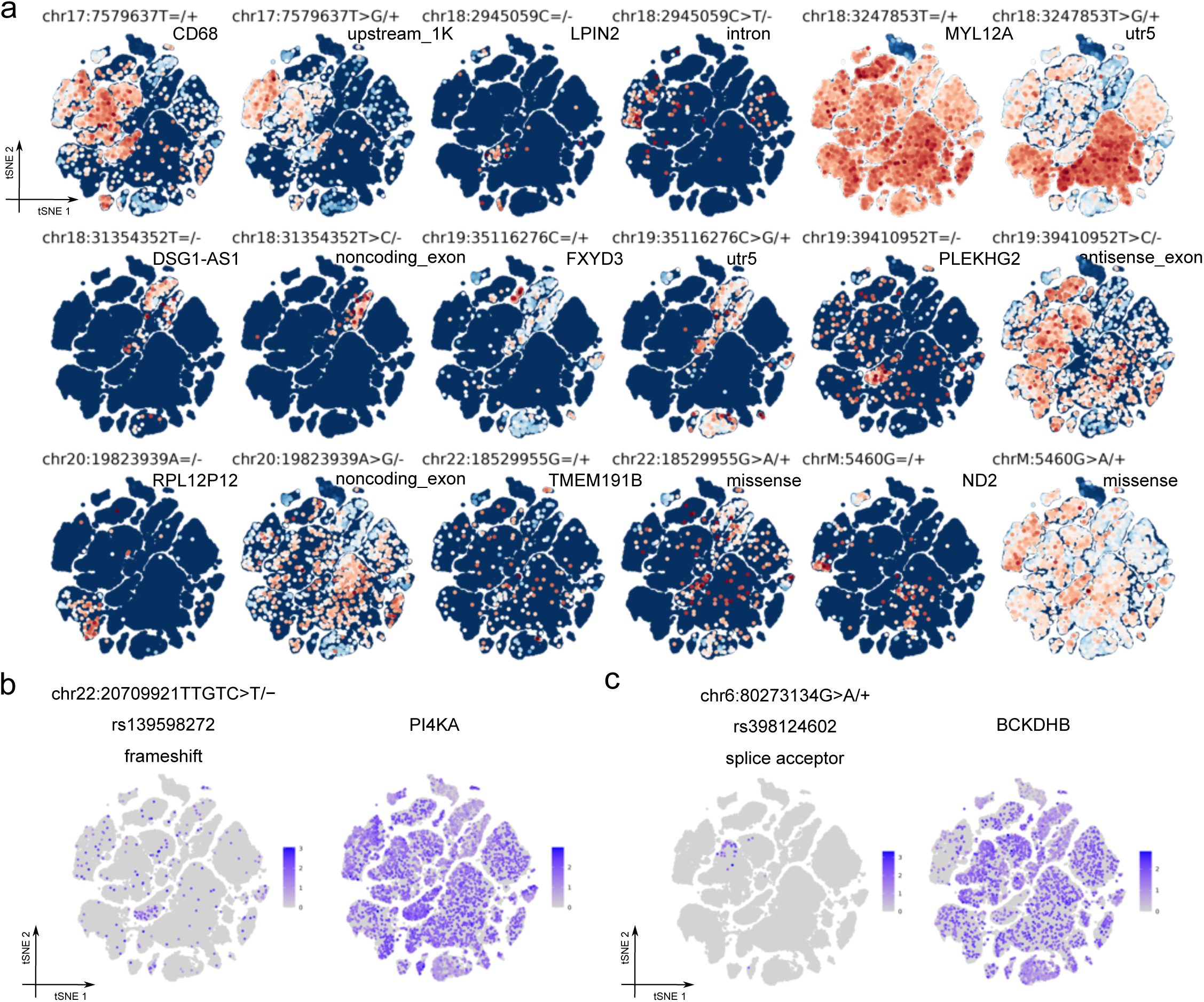
| Cases of allele-specific gene expression and selected genetic variants. Related to Figure 6. (a) Continuation of Supplementary Fig. 9. (b) tSNE visualization showing the normalized expression levels of EAT chr22:20709921TTGTC>T/-, a frameshift mutation (left) and its corresponding gene expression (right). (c) tSNE visualization showing the normalized expression levels of EAT chr6:80273134G>A/+, a splice acceptor mutation (left) and its corresponding gene expression (right).

